# The small ORF (smORF)-encoded polypeptide MIRINE promotes neuronal differentiation in vertebrate brain development

**DOI:** 10.64898/2026.10.31.762741

**Authors:** Yeji Kim, Benedikt Nilges, Ahmed Elhelbawi, Nathalie Krusy, Rościsław Krutyhołowa, Hannes C.A. Drexler, Laurent Nguyen, Sebastian A. Leidel

**Affiliations:** Department of Chemistry, Biochemistry and Pharmaceutical Sciences (DCBP), University of Bern, 3012 Bern, Switzerland; Max Planck Institute for Molecular Biomedicine, 48149 Muenster, Germany; Laboratory of Molecular Regulation of Neurogenesis, GIGA Institute, University of Liège, 4000 Liège, Belgium; Institute of Molecular Biology and Biophysics, ETH Zurich, 8093 Zurich, Switzerland; WELBIO department, WEL Research Institute, 1300 Wavre, Belgium

**Keywords:** SEPs, microprotein, neurodevelopment, ribosome profiling, small open reading frame

## Abstract

The vertebrate brain expresses an extensive repertoire of long-noncoding RNAs (lncRNAs), an emerging source of small open reading frame (smORF)-encoded polypeptides (SEPs) whose functions remain largely unknown. Using ribosome profiling and proteomics of zebrafish and mouse brain, we identified hundreds of translated smORFs. We characterized MIRINE, a SEP of 83 amino acids conserved across 205 vertebrate species. MIRINE contains a transmembrane domain that targets it to the plasma membrane and intracellular vesicles, and is enriched in cortical and hippocampal neurons, with expression rising during neurogenesis. Downregulation of *Mirine* increased cell proliferation and impaired neuronal maturation in neuroblastoma cells, in mouse primary cortical neurons and cortex, and in the zebrafish forebrain, identifying MIRINE as a regulator of the balance between neuronal proliferation and differentiation. Mechanistically, MIRINE associates with the synaptic-vesicle protein Synaptogyrin-3 and related membrane proteins. Our findings define a conserved, membrane-associated SEP controlling neuronal maturation and illustrate the functional potential of brain SEPs.

## Introduction

Proteins are the workhorses in every living cell. Their structure and function are determined by their amino acid sequence, which is encoded by an open reading frame (ORF) in their messenger RNA (mRNA). The identification and functional characterization of proteins is a key strategy in biology as it allows to pinpoint the functional elements within genomes that drive organismal physiology. Advances in functional genomics, guided by sophisticated computational methods, have greatly facilitated this approach and increased our understanding of biological processes at the cellular and organismal levels.

Over 20,300 protein coding genes are annotated in the human genome (Ensembl Genes 110 (GRCh38.p14)). However, genomics strategies have mostly focused on identifying proteins encoded by ORFs that are longer than 300 nucleotides and contain canonical start and stop codons (Basrai et al. 1997). In recent years, the discovery of atypical ORFs that do not match these parameters has sparked excitement (Andrea Pauli et al. 2015; Deutsch et al. 2026). These ORFs can be grouped by position on a transcript: (i) upstream ORFs (uORFs) in the 5’-untranslated regions of mRNA, (ii) downstream ORFs (dORFs) in the 3’-untranslated regions of mRNA, and (iii) small ORFs (smORFs) of less than 100 codons (Couso and Patraquim 2017). Many of these potential smORFs are located in uncharacterized RNAs that are annotated as long non-coding RNAs (lncRNAs). uORFs and dORFs function to post-transcriptionally regulate main ORFs and are being actively studied (Barbosa et al. 2013; Bazzini et al. 2014). However, identifying and characterizing smORFs and smORF-encoded polypeptides (SEPs) is still an emerging field. Identifying RNAs that code for smORFs is challenging because their size limits identification by genetics or computational approaches. An additional complication in identifying them is that smORFs do not always initiate translation at canonical ATG start codons (Ji et al. 2015). Therefore, transcripts containing smORF are often annotated as lncRNAs, and thousands of functional peptides may have evaded characterization, likely leaving various biological phenomena unexplained. Up to 40% of transcripts annotated as lncRNAs and pseudogene RNAs in human cells may therefore be translated into functional peptides (Ji et al. 2015).

A breakthrough for smORF discovery was the development of high-resolution ribosome profiling that allows for the identification of coding regions with high confidence. This strategy has been successfully applied to identify smORFs in different organisms (Aspden et al. 2014; Brar et al. 2012; Ingolia et al. 2011; Chen et al. 2020). However, aribosome profiling data alone does not prove the existence of a stable or physiologically relevant peptide (Brar and Weissman 2015). Therefore, it is crucial to establish the existence of SEPs at the protein level – for example with protein mass spectrometry. Advanced proteomics methods, such as shotgun proteomics (Alves et al. 2007), targeted proteomics (Michael A Gillette 2014), and human leukocyte antigen class I peptidomics (Bassani-Sternberg et al. 2015) have been used to confirm that candidate smORFs yield SEPs that are detectable in cells. For instance, an analysis of the proteomes of 80 human hearts by mass spectrometry identified 128 peptides from 86 SEPs encoded by transcripts previously considered non-coding (van Heesch et al. 2019). Systematic efforts to identify SEPs using translatome and proteome data have led to an increasing number of validated SEPs available in public repositories (Olexiouk et al. 2018; Hao et al. 2018). Nevertheless, the vast majority of putative smORFs likely await their verification at the protein level, and the cellular functions of SEPs remain unexplored.

Despite these challenges, multiple examples demonstrate the physiological importance of smORFs and SEPs in mitochondria (Bhatta et al. 2020; Makarewich et al. 2018; Stein et al. 2018; S. Zhang et al. 2020), development (Herberg et al. 2018; A. Pauli et al. 2014), cancer (T. Zhang et al. 2025; Kowar et al. 2025; Ouspenskaia et al. 2022), cellular stress (X. Li et al. 2025), and as neuropeptides (Fricker 2010; Guo et al. 2003; Pak et al. 2001). These examples highlight the high potential of smORFs and SEPs to uncover new and fascinating aspects of biology.

Due to the limited peptide space available to encode protein function in smORFs, we speculate that some SEPs may act in developmental signaling and in tissues such as the brain that are heavily reliant on these processes. The brain is one of the most complex organs, and many of its biological processes remain elusive. Identifying functional brain SEPs will deepen our understanding of biological mechanisms, especially since the brain is likely a rich source of SEPs, given its large number of expressed noncoding RNAs (Hezroni et al. 2015; Washietl et al. 2014).

Here, we describe MIRINE, a highly conserved SEP – also designated Sep18 – expressed during vertebrate brain development. In a ribosome profiling-based screen, we identified 198 and 121 smORF candidates in zebrafish and mouse brains, respectively. We selected candidates for further analysis based on proteomics, phylogenetic sequence conservation, and domain prediction. MIRINE (MIcRoprotein Inducing NEuronal differentiation) is an 83-amino-acid-long SEP that localizes to the plasma membrane and vesicles. We observed that *Mirine* is expressed during the later stages of neurogenesis and in mature neurons. Knocking down *Mirine* perturbs the homeostasis between the proliferation and differentiation of neuronal progenitors, restricting their maturation in cell culture and in mice. This regulation may be mediated by a physical association with Synaptogyrin-3 (SYNGR3). Our findings identify MIRINE as a new modulator of neuronal differentiation and shed light on the potential of SEPs in the underlying regulatory mechanisms of neuronal maturation during development.

## Materials and methods

### Cell culture and animal experiments

HEK293T cells and Neuro2a (N2A) cells were purchased from ATCC (VA, United States). Cells were cultured in Dulbecco’s modified Eagle’s medium (DMEM) supplemented with 10% Fetal Bovine Serum (FBS; Merck), 1% Penicillin-Streptomycin (Thermo Fisher, MA, United States), 1% Non-essential amino acids, and 1% GlutaMAX™ supplement (Thermo Fisher, MA, United States). Cells were passaged every 2-3 days at 80% confluency using 0.1% Trypsin-EDTA (Invitrogen, MA, United States). For the differentiation of N2A cells, the standard medium was replaced by DMEM containing 2% FBS and 20µM all-trans-retinoic acid (Sigma-Aldrich, MO, United States) for 48 h. Mice and zebrafish were used following Belgian, German, and European animal protection guidelines. All procedures were ethics-approved by the University of Liège animal ethic committee (#14-1638) and the Landesamt für Natur, Umwelt und Verbraucherschutz (LANUV) of the state of North Rhine-Westphalia, Germany (#84-02.05.20.13.001), and supervised by authorized staff. Mice were bred in-house and maintained at 19–22 °C, 40–50% humidity, on a reversed 12:12 light/dark cycle (lights on at 23:00), with ad libitum food and water.

### Ribosome profiling

Ribosome profiling of adult zebrafish (AB) and mouse (C57BL6) brains was performed based on a published protocol (Y. Kim et al. 2021; Ingolia et al. 2009; Lecanda et al. 2016; Sharma et al. 2021) with modifications. Briefly, brains were isolated from sacrificed animals and snap-frozen in liquid nitrogen until further processing. One hemisphere of a mouse brain or 6-10 pooled zebrafish brains were washed with ice-cold phosphate-buffered saline (PBS) containing 100 μg/ml Cycloheximide (CHX) and lysed in lysis buffer (50 mM Tris-HCl pH 7.5, 12 mM MgCl2, 50 mM NH4Cl, 2 mM DTT and 100 μg/ml CHX). Lysates were cleared by centrifugation at 12,000x g and 4 °C for 5 min. 10 A260 units of the supernatant were treated with 800 U RNase I (Thermo Fisher Scientific, MA, United States) for 10 min at 25 °C. The reaction was stopped by the addition of 15 μl SuperaseIn (Thermo Fisher), and the monosome fraction was isolated from a 10-50% (w/v) sucrose gradient. 10 μg of RNA of the monosome fraction were separated on a 15% polyacrylamide gel containing 8 M Urea. 28-30 nt long footprints were excised. rRNAs were depleted using the Ribo-Zero Gold rRNA Removal Kit (Illumina, CA, United States) and biotinylated DNA oligos. The libraries were sequenced with a HiScanSQ (Illumina, CA, United States) generating 50 nt single-end reads.

### RNA-sequencing

To isolate total RNA from zebrafish and mouse brains, snap-frozen brains were thawed in 1 ml of TRI Reagent (Sigma-Aldrich, MO, United States) and 5x vortexed with glass beads for 10 s with intermittent cooling on ice for 15 s. The lysates were 10x triturated through a 20G needle and incubated for 5 min at room temperature (RT). For total RNA isolation from N2A cells, cells were harvested mechanically at 70-80% confluency. RNA was isolated using TRI Reagent (Sigma-Aldrich, MO, United States) according to the manufacturer’s instructions. Residual genomic DNA was removed by incubating RNA with 0.2 U TURBO DNase (Invitrogen by Thermo Fisher Scientific, MA, United States) per 1 μg of total RNA for 30 min at 37 °C on a shaker. Samples were qualified with a Bioanalyzer 2100 (Agilent Technologies, CA, United States) and only samples with RNA Integrity Index values >8 were used. Sequencing libraries were prepared using a TruSeq Stranded mRNA Library Prep Kit (Illumina, CA, United States).

### Analysis of ribosome profiling and RNA-sequencing

For ribosome profiling libraries, adaptor sequences and four random nucleotides were removed before alignment by using fastx-toolkit (Gordon 2010). Libraries were mapped to zebrafish or mouse rRNA, and the remaining reads were aligned to GRCz10 (zebrafish) or GRCm38 (mouse) using bowtie (Langmead et al. 2009). Differential gene expression was analyzed by DESeq2 (Love et al. 2014) and gene ontology (GO) term analysis was performed using the TopGO R package (Alexa and Rahnenführer 2026).

### ORF discovery

An extended zebrafish transcriptome was generated to identify novel ORFs. Ribosome profiling and RNA-seq libraries were aligned to the danRer10 genome using the splice-aware aligner TopHat2 (version 2.0.11) (Kim et al. 2013) supplied with transcript models from the ensGene track of the UCSC Genome Browser. Splice junctions found during this run were also used during the mapping of data for ORF identification. Mapped reads were used to assemble transcripts with Cufflinks (version 2.0.2) (Trapnell et al. 2010) using the ensGene annotation to identify known transcripts, merging elements separated by less than 200 nucleotides. RiboTaper (Calviello et al. 2015) required transcript sequences for ORF finding, based on the danRer10 genome alongside a transcriptome annotation. The transcriptome used for this purpose was the one generated above, merged with coding and noncoding ENSEMBL genes (ensGene) as well as published lincRNAs (Kaushik et al. 2013; Ulitsky et al. 2011; Andrea Pauli et al. 2012), which was sorted by position in the genome using BEDTools sort (version 2.0.17) (Quinlan and Hall 2010). The information was transferred into RiboTaper-readable annotation files. Footprints of 29, 30 or 31 nt were chosen to determine ORFs, with the P-site location defined as 12 nt from the start of reads (Supplementary Figure 1A and B).

### Plasmid generation

miRFP670nano-pcDNA was a gift from V. Verkhusha (Addgene plasmid # 127443). The full sequence of *Mirine* or transmembrane domain-truncated *Mirine* was PCR-generated and subcloned at the C-terminus of miRFP670nano. Knockdown constructs for *in utero* electroporation were subcloned into pSilCAEGFP (Addgene plasmid # 52675) using NEBuilder® HiFi DNA Assembly (New England Biolabs, MA, USA). Endotoxin-free constructs were isolated using PureLink™ HiPure Expi Plasmid Megaprep Kit (Invitrogen™ by ThermoFisher Scientific, MA, USA). shRNA targeting *Mirine* are as follows: GCTATGGATTGCCATCATA (shRNA#1); GGACTCACCTAGTAAGGTT (shRNA#2).

### Plasmid transfection

Overexpression constructs were transfected using FuGENE^®^ HD Transfection Reagent (Promega, WI, United States) with a ratio of 1:3.5 of DNA:Reagent according to the manufacturer’s instruction. Transfected cells were analyzed at 48-72 h following transfection.

### siRNA Knockdown

Scrambled control, *Mirine*, and *Syngr3* SiPOOL oligos were purchased from SiTOOLs Biotech GmbH (Martinsried, Germany). SiRNAs were transfected into N2A cells using Lipofectamine™ RNAiMAX Transfection Reagent (ThermoFisher Scientific, MA, USA) according to the manufacturer’s instructions. Transfected cells were harvested after 72 h for subsequent analysis.

### Morpholino experiments

A translation-blocking morpholino (MO) (Gene Tools, OR, USA) was injected into one-cell-stage embryos at a concentration of 300 μM (500 pl per embryo). As a negative control, Gene Tools non-targeting standard MO was used. The embryos were imaged using a Nikon SMZ18 stereo microscope with a dorsal orientation. The diameter of the eye, forebrain and midbrain of the MO-injected embryos was measured in Fiji at 5 dpf and normalized to body length. Morpholino sequence: CGCAGCCATAAATGACAGATCCATG.

### Western blot

Cells were lysed in RIPA buffer after washing with ice-cold PBS. Lysates were denatured by boiling at 95 °C for 5 min or incubating at 37 °C for 15 min (for Tricine-SDS-PAGE) with NuPAGE^™^ LDS Sample Buffer and sample-reducing reagent (ThermoFisher Scientific, MA, USA). Protein lysates were separated on acrylamide gels of varying percentages based on the target protein size: 10% and 15% SDS-PAGE gel, 16% Tricine-SDS gel, or mPAGE 4-12% Bis-Tris Precast Gel (Sigma-Aldrich, MO, United States). Gels were transferred either by semi-dry transfer at 10 V for 20 min for proteins smaller than 20 kDa, or by wet transfer at 50 V for 2 h for larger proteins. PVDF membranes were blocked with 4% non-fat milk in TBS for 1 h at RT and incubated with designated primary antibodies at 4 °C overnight. HRP-conjugated or fluorescent secondary antibodies were used for detection. Membranes were scanned using an Azure Sapphire Biomolecular Imager (Azure Biosystems, CA, USA). Quantifications were performed using Azurespot (Azure Biosystems, CA, USA) and FIJI. Antibody concentrations and conditions are listed in Table 1.

**Table 1.** Antibody list.

| Antibody | Product number | Species | Dilution | Experiment |
|---|---|---|---|---|
| MIRINE | Custom antibody | Rabbit | 1:100 | WB |
| GFP | Nacalai Tesque GF090R | Rat | 1:500 | IHC |
| COPG | Sc-393977 | Mouse | 1:1,000 | WB |
| TUBB2 | Sc-47751 | Mouse | 1:1,000 | WB |
| TUBB3 | Sc-51670 | Mouse | 1:1,000 | WB |
| SYNGR3 | Sc-514081 | Mouse | 1:1,000 | WB, IHC |
| ARF1 | Sc-53168 | Mouse | 1:1,000 | WB |
| β-actin | GeneTex 44363 | Rabbit | 1:2,000 | WB |
| FLAG-M2 | Sigma 422303 | Mouse | 1:2,000 | WB |
| Rat IgG (Alexa 488) | Jackson ImmunoResearch 712-545-153 | Donkey | 1:500 | IHC |
| Mouse IgG (AzureSpectra 800) | Azure AC2135 | Goat | 1:25,000 | WB |
| Mouse IgG (AzureSpectra 700) | Azure AC2129 | Goat | 1:25,000 | WB |
| Rabbit IgG (AzureSpectra 700) | Azure AC2128 | Goat | 1:25,000 | WB |
| Rabbit IgG (AzureSpectra 800) | Azure AC2134 | Goat | 1:25,000 | WB |
| Rabbit (HRP) | AC2114 | Goat | 1:25,000 | WB |

### Peptide antibody generation

Peptides were synthesized and used for antibody generation by Eurogentec (Seraing, Belgium) using the standard immunization program for peptide antibodies.

### qRT-PCR

Total RNA was isolated from N2A cells using TRI Reagent (Sigma-Aldrich, MO, United States) according to the manufacturer’s instructions. cDNA was synthesized from 1 μg of RNA using GoScript^™^ Reverse Transcription System (Promega, WI, United States) with random priming. qRT-PCR mixtures were prepared with KAPA SYBR FAST qPCR mix (Sigma-Aldrich, MO, United States) and amplified on a Rotor-Gene Q (Qiagen, Hilden, Germany). For comparative analyses, target gene expression was normalized to mouse U1 snRNA or *Rpl18a*. Primers used in qRT-PCR are listed in Table 2.

**Table 2.**
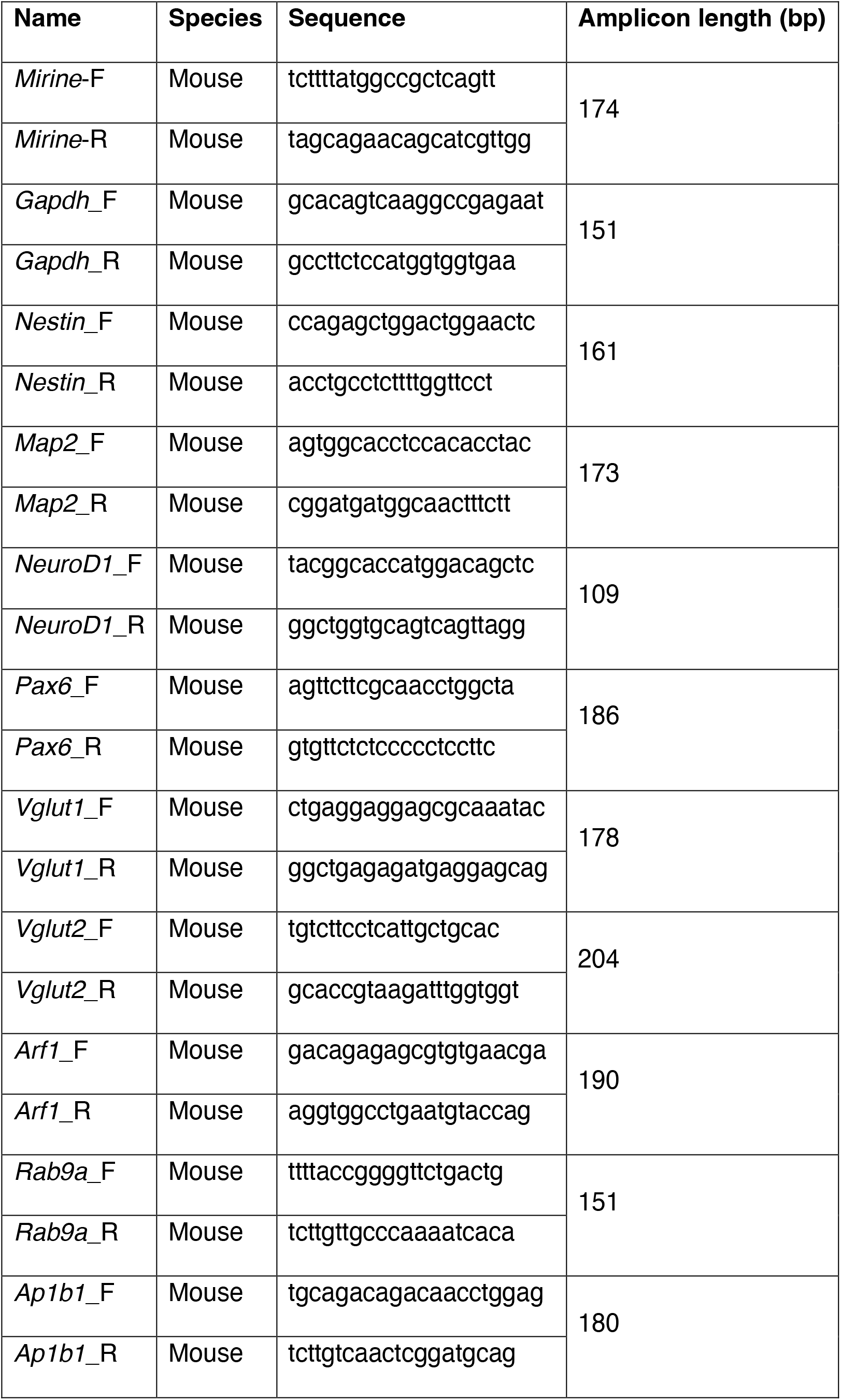
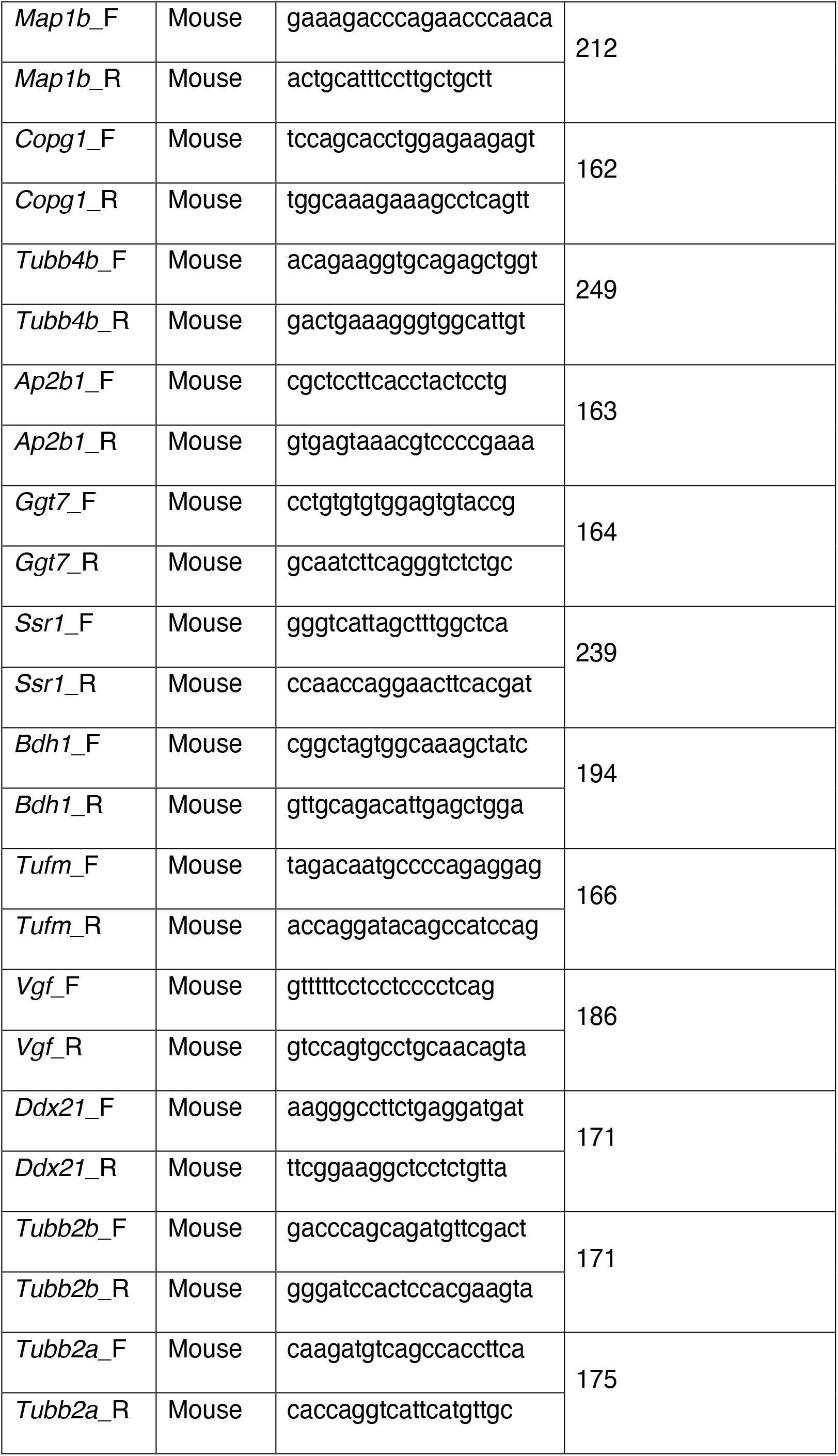
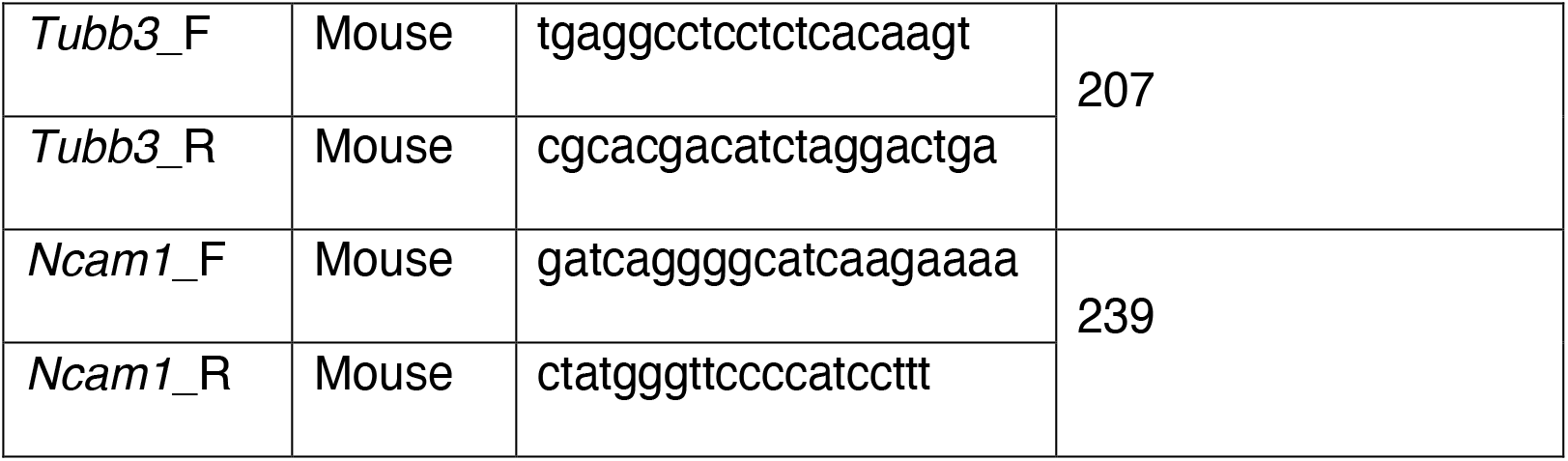
Primer list.

### *In situ* hybridization (ISH)

*In situ* probes were derived from PCR-amplified cDNA fragments of zebrafish or mouse. For zebrafish, ABxTL zebrafish embryos were fixed at designated developmental time in 4% paraformaldehyde. Rehydrated embryos were permeabilized by Proteinase K (AppliChem) and hybridized with 2 μg/ml RNA probes at 65 °C overnight. For mouse, ISH was performed as described before (Cau et al. 1997; Wilkinson and Nieto 1993). Primers used for generating probes are listed in Table 3.

**Table 3.**
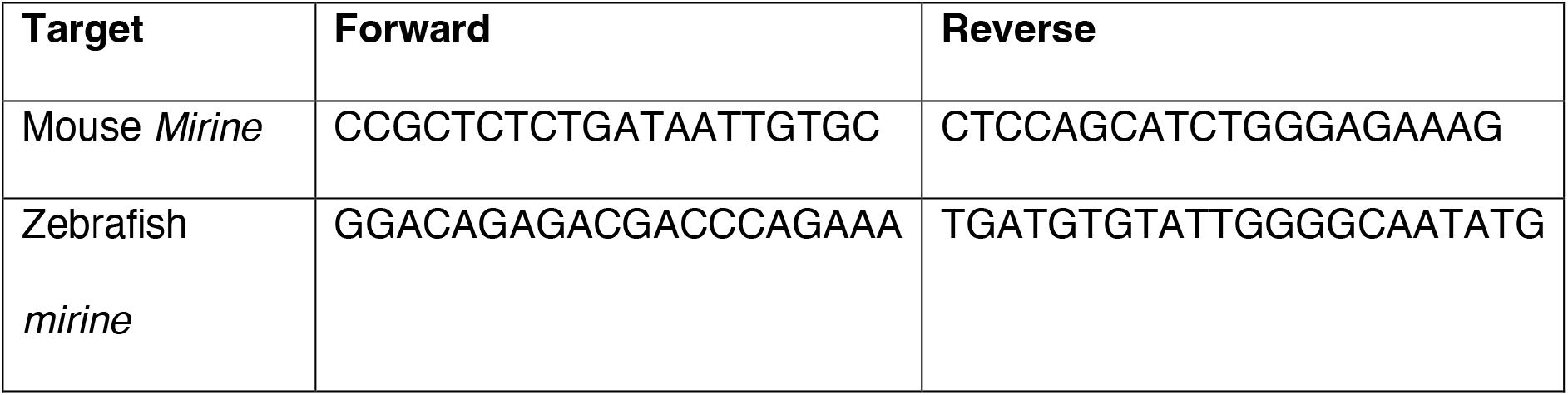
Primers used for generating ISH probes.

### Immunocytochemistry

HEK293T cells and N2A cells were fixed using fresh 10% Formalin (Sigma-Aldrich, MO, United States). Fixed cells were blocked with 5% donkey serum (adding 0.2% Triton X-100 if permeabilization was needed) and incubated using primary antibodies at 4 °C overnight. Alexa 488 or 594-conjugated secondary antibodies were bound for 1 h at RT. Cells were counter-stained with DAPI and CellMask™ Green Plasma Membrane Stain (C37608, Thermo Fisher Scientific) for subcellular organelle visualization. Cells were imaged using CSU-X1 (Nikon, Japan). Indicated images were rendered using Huygens software.

### Tissue processing for immunohistochemistry

Zebrafish brains and mouse embryonic brains were dissected in ice-cold PBS and fixed at 4 °C in 4% paraformaldehyde (PFA) for 20 min (zebrafish) or 1 h (mouse). Fixed zebrafish brains were embedded in paraffin after dehydration by ethanol and xylene. Fixed mouse brains were cryoprotected overnight in 20% sucrose in PBS at 4 °C, embedded in OCT Compound (VWR International, Leuven, Belgium) and sectioned into 12-16 μm slices onto glass slides (SuperFrost Plus, VWR International, Leuven, Belgium) using a cryostat.

### Immunohistochemistry

Frozen cryostat sections were washed three times in PBS-Triton 0.1% (PBST) and blocked for 1 h at room temperature in PBST containing 10% donkey serum (Jackson ImmunoResearch Laboratories, PA, USA). Sections were incubated overnight at 4 °C with the primary antibodies. Pretreatment with 2 N HCl for 30 min or with Target Retrieval Solution (Agilent Technologies, CA, United States) for 20 min at 95 °C prior to incubation with primary antibody was required for detection of fluorescence. After washing, sections were incubated for 1 h at RT with secondary antibodies coupled to Alexa-488, Alexa-647 or Alexa-555 (Jackson ImmunoResearch Laboratories or Life Technologies). Nuclei were counterstained with Hoechst 33342 (1:1,000, Sigma-Aldrich, MO, United States) or DAPI (1:10,000, Sigma-Aldrich, MO, United States), washed in PBST and mounted using Mowiol Mounting Medium (Sigma-Aldrich, MO, United States). Slides were stored in the dark at 4 °C. For image analyses, sections were imaged using a A1Ti confocal microscope (Nikon, Tokyo, Japan) and FIJI software. The list of antibodies used in the staining were indicated in Table 1.

### Immunoprecipitation-mass spectrometry (IP-MS)

N2A cells were harvested 48 h post-transfection. 50 μL of Dynabeads protein G (Thermo Fisher Scientific, CA, United States) was coupled to 1 μg anti-FLAG M2 antibody (F3165, Sigma-Aldrich, MO, United States) or anti-Mouse IgG antibody (Santa Cruz Biotechnology, TX, United States) for 30 min at RT. A minimum of 2 mg of cell lysates were incubated with antibody-coupled beads for 30 min at RT with agitation. Antigen-antibody complexes were washed 3x with wash buffer (10 mM Tris-HCl pH 7.4, 1 mM EDTA, 1 mM EGTA pH 8.0, 150 mM NaCl, and 1% Triton X-100), 3x with PBS + 0.2% Tween 20, and 2x with PBS. Samples were eluted in 50 mM glycine pH 2.7 with NuPAGE LDS Sample Buffer and reducing reagent (ThermoFisher Scientific, MA, USA) at 70 °C for 10 min and shortly run on 15% SDS-PAGE gel for excision. Proteins were digested by Chymotrypsin + Glu-C 1:1 mix at a concentration of 10 ng/μL. An aliquot of 5 μL of each digest was then analyzed on a nano-liquid chromatography system coupled with tandem mass spectrometry consisting of an Ultimate 3000 UPLC and an orbitrap tribrid LUMOS (ThermoFisher Scientific). Samples were loaded onto a pre-column (C18 PepMap 100, 5 μm, 100 A, 300 μm i.d. x 5 mm length) at a flow rate of 10 μl/min with solvent C (0.05% TFA in water/acetonitrile 98:2). After loading, peptides were eluted in backflush mode onto a homemade C18 CSH Waters column (1.7 μm, 130 Å, 75 μm x 20 cm) by applying a 40-min gradient of 5% acetonitrile to 40% in water, 0.1% formic acid, at a flow rate of 210 nl/min. Data were acquired in data dependent mode with precursor ion scans recorded in the orbitrap at a resolution of 120′000 (at m/z = 250) parallel to top speed fragment spectra of the most intense precursor ions in the linear ion trap for a cycle time of 3 s maximum, applying a 30-s exclusion time window.

MS data was analyzed with MaxQuant, version 1.6.14.0 (Cox and Mann 2008) against a database of concatenated protein sequences from SwissProt Mus Musculus (version 2022_01), common contaminants, and the reversed decoy sequences. Search parameters included a mass tolerance of 10 ppm and 0.4 Da at MS1 and MS2 level; static carbamidomethyl on cysteine, and variable oxidation on methionine and acetylation of protein N-termini; combined chymotrypsin and GluC cleavage specificity; and 1% FDR filtering on PSM, peptide, and protein level in addition of accepting protein groups only when identified by two peptides. Proteins identified in at least three replicates with significance were considered as interacting partners.

### *In utero* electroporation

*In utero* electroporation was performed as previously described (Creppe et al. 2009; Nguyen et al. 2006) with minor modifications. Briefly, uteri of timed-pregnant mothers (14.5 days) anaesthetized with isoflurane in an oxygen carrier (Abbot Laboratories Ltd, Kent, UK) were exposed through a 1.5 cm incision in the ventral peritoneum. Embryos were carefully lifted using ring forceps through the incision and placed on humidified gauze pads. Plasmid DNA solution (2 μg/μl) were mixed with 0.05% Fast Green (Sigma Aldrich, MO, United States) for injection. The plasmid was injected through the uterine wall into the telencephalic vesicle using pulled borosilicate needles and a Femtojet microinjector (VWR International, Leuven, Belgium). Five electrical pulses were applied at 35 V (50 ms duration) across the uterine wall at one-second intervals using 5 mm platinum tweezers electrodes (CUY650P5, Sonidel Limited, Dublin, Ireland) and an ECM-830 BTX square wave electroporator (VWR International, Leuven, Belgium). The uterine horns were then replaced in the abdominal cavity and the abdomen walls and skin were sutured using a surgical needle and thread. The pregnant mouse was injected with buprenorphine (Temgesic®, Schering-Plough, Brussels, Belgium) and warmed on a heating pad until it woke up. The whole procedure was performed within 45 min. Pregnant mice were sacrificed by neck dislocation at E15.5 (for primary neuron culture), E16.5 (EdU incorporation analysis) or P2 (migration analysis) and embryos were processed for tissue analyses to study radial migration.

### EdU incorporation analysis

EdU incorporation analysis was performed using a Click-iT™ EdU Cell Proliferation Kit for Imaging (C10340, ThermoFisher Scientific) or a Click-iT™ Plus EdU Pacific Blue™ Flow Cytometry Assay Kit (C10636, ThermoFisher Scientific) according to the manufacturer’s instruction. Briefly, N2A cells were plated with either control or si*Mirine* for 48 h. 2h prior to harvesting, EdU was added to the cells. Cells were co-stained with propidium iodide and analyzed by ImageStreamX MkII. For mouse *in vivo* experiments, electroporated mice were injected EdU with IP injection 1 h or 24 h before sacrifice.

### Mouse primary cortical neuron culture

Scrambled or *Mirine* knockdown construct-injected embryonic mice were dissected on E15.5. GFP signal-positive cortices were dissociated by 0.05% Trypsin for 20 min at 37 °C. 20,000 cells were plated on a 24-well plate and cultured in Neurobasal™ Medium (Thermo Fisher, MA, United States) supplemented with 2% B27, 1% GlutaMAX and 1% Penicillin-Streptomycin (Pen-Strep) for 7-9 days. Media were changed every 3 days. Cells were fixed in 4% PFA and proceeded to immunofluorescent staining.

### Image analysis and statistics

Fluorescent images were quantified by counting a minimum of 3 brains from each group. Magnified fields were acquired on 2-3 adjacent sections with confocal microscopy. At least three brains were analyzed for each experimental group. A Sholl analysis of primary cortical neurons was performed using the SNT plugin of FIJI. Statistics for dual comparisons were performed using unpaired Student’s t-test, and multiple comparisons were performed using one- or two-way ANOVA followed by a post hoc test.

### AlphaFold3 modeling of MIRINE–Synaptogyrin-3 complexes

Sequences of MIRINE/Mirine from zebrafish (*Danio rerio*; UniProt A0A286YBL2), mouse (*Mus musculus*; J3QPM6) and human (*Homo sapiens*; Q9NPU4), and of the corresponding Synaptogyrin-3 orthologs (zebrafish Synaptogyrin-3a, Q503N6; mouse, Q8R191; human, O43761), were retrieved from UniProt. Each MIRINE-Synaptogyrin-3 pair was modeled as a 1:1 heterodimer using the AlphaFold Server (AlphaFold3; Abramson et al. 2024) with default settings and an automatically assigned random seed. Top scoring models were inspected and visualized in UCSF ChimeraX (Meng et al. 2023). PAE plots were generated using a custom Python script.

## Results

### Ribosome profiling identifies MIRINE, a brain-enriched SEP

We sought to identify new functional SEPs in the vertebrate brain. To this end, we performed ribosome profiling and RNA-seq using brains from adult mice and zebrafish. To exclude false positives and accurately identify true translation events, it is essential to generate sequencing libraries displaying robust 3-nt periodicity. Our ribosome profiling data revealed striking periodicity for footprint sizes ranging from 29 nt to 31 nt, allowing us to identify potential SEPs (Supplementary Figure 1A and B). To discover uncharacterized smORFs, we mapped our reads exclusively to putative lncRNAs and employed RiboTaper to identify unannotated smORFs (Calviello et al. 2015). SmORFs are defined as small ORFs that are (i) confirmed by ribosome profiling and RNA-seq, (ii) supported by a significant number of in-frame reads, and that (iii) contain a start and a stop codon. We identified a total of 6967 and 2884 smORF candidates in lncRNAs of zebrafish and mice respectively. Next, we categorized these smORFs based on their relative position on the transcript. The vast majority of smORFs (68% in zebrafish; 59% in mice) were uORFs, followed by ORFs on uncharacterized or lncRNA transcripts (ncORFs) (29% in zebrafish; 34% in mice). We detected only a small number of dORFs (2% in zebrafish; 7% in mice) (Supplementary Figure 1C). The detected smORFs were on average significantly shorter (54 nt in zebrafish; 117 nt in mice) than canonical ORFs (1194 and 1170 nt, Supplementary Figure 1D). To identify uncharacterized SEPs, we excluded uORFs and dORFs since they mostly perform regulatory roles for the translation of their main ORFs. First, we removed smORFs that matched known mRNA by BLAST, leaving 198 (zebrafish) and 121 (mice) smORF candidates (Supplementary Figure 1C, Supplementary Table 1 and 2). Second, to select SEP candidates with high functional potential, we applied several metrics: (i) detection in in-house shotgun proteomics data, (ii) phylogenetic conservation (Lin et al. 2011), and (iii) predicted domains (Petersen et al. 2011; Krogh et al. 2001; Letunic et al. 2015; Schultz et al. 2000) (Supplementary Figure 1E).

Based on this metric, we selected a putative SEP (ENSDART00000152821.1 in zebrafish; *D430019H16Rik* in mice; *C14orf132* in human) based on its high phylogenetic conservation in vertebrates and domain prediction (Figure 1A). Later, we named it MIRINE (MIcRoprotein Inducing NEuronal differentiation). Of note, MIRINE has been characterized under the designation SEP18 in vascular smooth muscle cells (Li et al. 2023). Its protein sequence is highly conserved in 205 vertebrate species including primates, rodents, and fishes (Figure 1B) and its lncRNA is expressed in a tissue-specific manner, predominantly in the brain (Supplementary Figure 2A). To determine its cellular localization, we generated peptide antibodies directed against zebrafish Mirine. We confirmed its expression in zebrafish brain by western blot and showed specificity by peptide competition (Figure 1C). Furthermore, GTEx data showed that the human orthologue of the *MIRINE* transcript is predominantly expressed in brain, but also shows weak expression in lung, pituitary gland, and esophagus (Supplementary Figure 2A), a pattern, similar to the mouse and zebrafish transcripts. We used AlphaFold2 to predict MIRINE’s structure and functional domains (Supplementary Figure 2B) (Jumper et al. 2021; Varadi et al. 2022). Its C-terminus is hydrophobic and likely contains a transmembrane domain. To test its subcellular localization, we fused MIRINE to the small miRFPnano670 tag (Oliinyk et al. 2019). Strikingly, miRFPnano670-MIRINE localizes to the plasma membrane and organellar membranes but loses its membrane localization when the putative transmembrane domain is truncated, supporting a functional role of the predicted transmembrane domain (Figure 1D). Next, we analyzed the expression patterns of *Mirine* during development in zebrafish, mice and humans using *in situ* hybridization and public RNA-seq data. In zebrafish, the *mirine* transcript is expressed globally early during development. However, at later stages, the expression of *mirine* was restricted to the head and spinal cord (Figure 1E). In mouse, we observed that *Mirine* mRNA levels increase during brain development with a peak of expression at E14.5 while maintaining lower levels in tissues such as heart, lung, kidney, liver, stomach, and intestine (Supplementary Figure 2C). Taken together, we identified MIRINE, a SEP conserved in vertebrates, which is predominantly expressed in the vertebrate brain with increasing expression during development.

**Figure 1.**
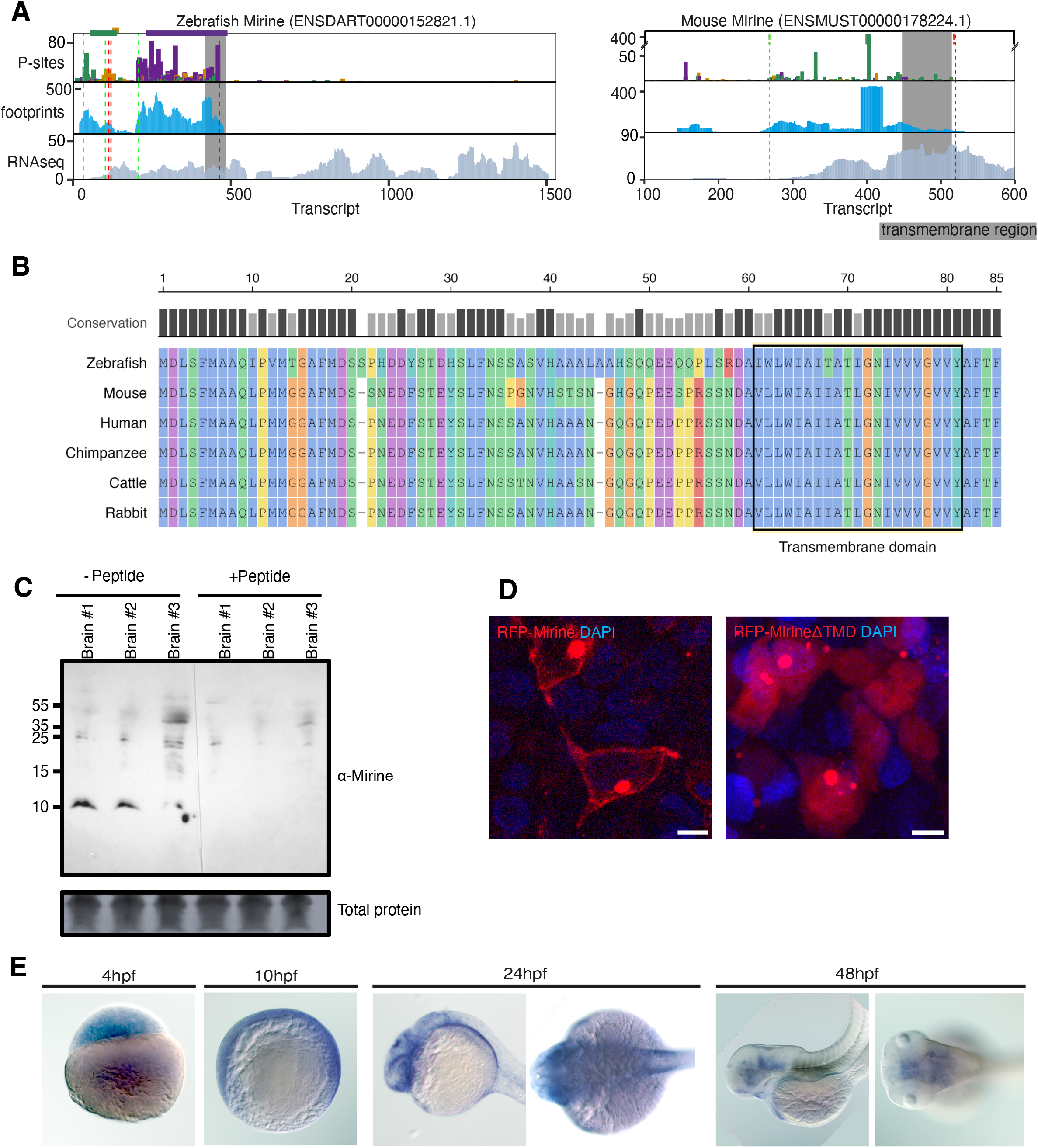
Expression of MIRINE in the vertebrate brain. (A) Visualization of *mirine*/*Mirine* translation in the zebrafish (left) and mouse (right) brain by ribosome profiling. (B) Protein sequence conservation across selected vertebrate species. (C) Western blot analysis of MIRINE protein expression in the zebrafish brain. (D) Subcellular localization of miRFP670nano-MIRINE (red) in HEK293T cell line. Nuclei are stained by DAPI (blue). (E) *In situ* hybridization of *mirine* mRNA during zebrafish embryonic development.

### *Mirine* is expressed in developing neurons

Next, we investigated the expression of *Mirine* in the brain in more detail. While zebrafish allowed initial validation of expression and localization, we used mice as our main model to dissect neurogenesis at single-cell resolution, given the more detailed and comprehensive cell-type annotation resources available. We employed the public scRNA-seq data of developing mouse brain (La Manno et al. 2021) to explore the expression pattern of *Mirine* to diverse brain regions and cell types. At E14.5 embryonic mouse brain, *Mirine* is predominantly expressed in neurons but relatively low in the non-neuronal cells (Figure 2A). As we observed that *Mirine* was highest in neurons followed by neuroblasts, we compared the expression levels of *Mirine* in developing stages of neurons (Radial glia, neuroblasts, and neurons) during embryonic development. In the time-course analysis, we observed that *Mirine* mRNA expression is increased around E10-E11, when neuronal differentiation starts (Figure 2B). Also, amongst all annotated cell types, neurons express the highest *Mirine* during development, which suggests its function in neuronal maturation (Supplementary Figure 3A-B). Next, we investigated whether *Mirine* is expressed in specific types of neurons in the brain. We compared the expression levels of *Mirine* between glutamatergic neurons and GABAergic neurons and found that *Mirine* was relatively highly expressed in glutamatergic neurons (Supplementary Figure 3C). In particular, in the forebrain, the expression levels between glutamatergic and GABAergic neurons showed significant differences (Figure 2C, p=0.008). In addition, using *in situ* hybridization, we confirmed that *Mirine* mRNA is enriched in the cortex of embryos and adult mice (Figure 2D and Supplementary Figure 3D). This expression pattern suggests that *Mirine* expression increases during neuronal differentiation. This is also reflected in the public bulk RNA-seq time course we analyzed (Supplementary Figure 2C) (Sloan et al. 2016). This indicates that *Mirine* is expressed during neuronal maturation.

**Figure 2.**
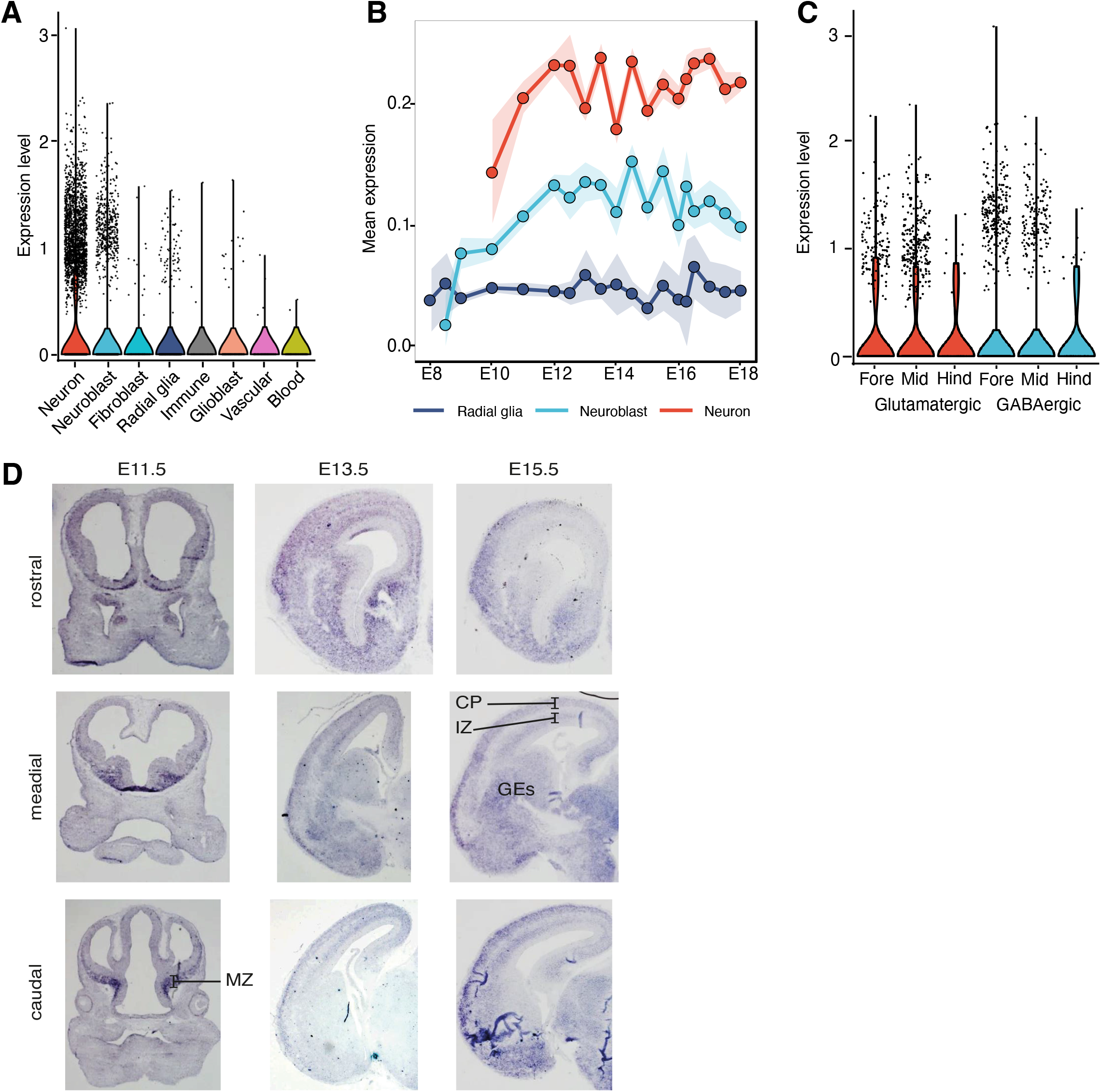
Neuron-enriched expression of *Mirine* in the mouse brain. (A) RNA expression of *Mirine* in the E14.5 mouse brain. (B) RNA expression of *Mirine* during embryonic mouse brain development in neuronal cells. (C) RNA expression of *Mirine* in E14.5 mouse brain subregion neurons. (D) *In situ* hybridization of *Mirine* during mouse brain development (E11.5-E15.5); cortical plates (CP); intermediate zone (IZ); ganglionic eminence (GE); marginal zone (MZ). scRNA-seq resource from La Manno et al. 2021 (A-C).

### Depletion of MIRINE shifts cells towards proliferation

To decipher the function of MIRINE, we knocked down its mRNA in mouse neuroblastoma N2A cells using siRNAs (Figure 3A). Strikingly, upon *Mirine* knockdown, N2A cells grew faster than control cells (Figure 3B). We verified the increased proliferation rates of knockdown *Mirine* cells by EdU incorporation and performed a cell cycle analysis by flow cytometry (Figure 3C and 3D). The number of EdU-positive cells increased in response to *Mirine* downregulation (Figure 3C). Furthermore, the number of cells in S phase was larger in the *Mirine* knockdown (Figure 3D). Therefore, we harvested cells 72 h after transfection and performed RNA-seq of control cells and si*Mirine* cells to identify transcripts that are differentially expressed upon *Mirine* knockdown. We observed that genes implicated in cell division and proliferation were strikingly upregulated (Figure 3E; red genes), while genes enriched in neuronal functions such as neurotransmitters and synapses were downregulated (Figure 3E; blue genes). Consistently, Gene Ontology (GO) terms related to proliferation and cell division were enriched in upregulated genes, while neuronal terms were enriched in downregulated genes (Supplementary Figure 4A). This indicates that in the absence of MIRINE, cells proliferate faster at the expense of differentiation. To test whether depletion of *Mirine* affects neuronal differentiation, we starved N2A cells to induce spontaneous differentiation. Indeed, *Mirine* depleted cells showed thicker neurites with fewer neurite branches, indicating that they are less differentiated (Figure 3F). To confirm the effect of MIRINE in neuronal differentiation, we used an orthogonal model and treated N2A cells with 20 µM of all-trans-retinoic acid (RA) for 2 days and compared the expression profile of si*Mirine* and wt cells by RNA-seq. In differentiated wild-type cells, *Mirine* expression was upregulated compared to the undifferentiated cells (Supplementary Figure 4B). However, not all neuronal marker genes were downregulated in *Mirine* knockdown cells. While neuronal maturation markers, *Tubb3* and *Tbr1* were reduced in *Mirine* knockdown cells, genes like *Ascl1*, *Map2*, *Nes*, and *Pax6* were expressed at relatively high levels (Figure 3G). Collectively, these data indicate that MIRINE functions as a regulator that restrains proliferation and supports neuronal differentiation, and its depletion shifts cells toward a proliferative state.

**Figure 3.**
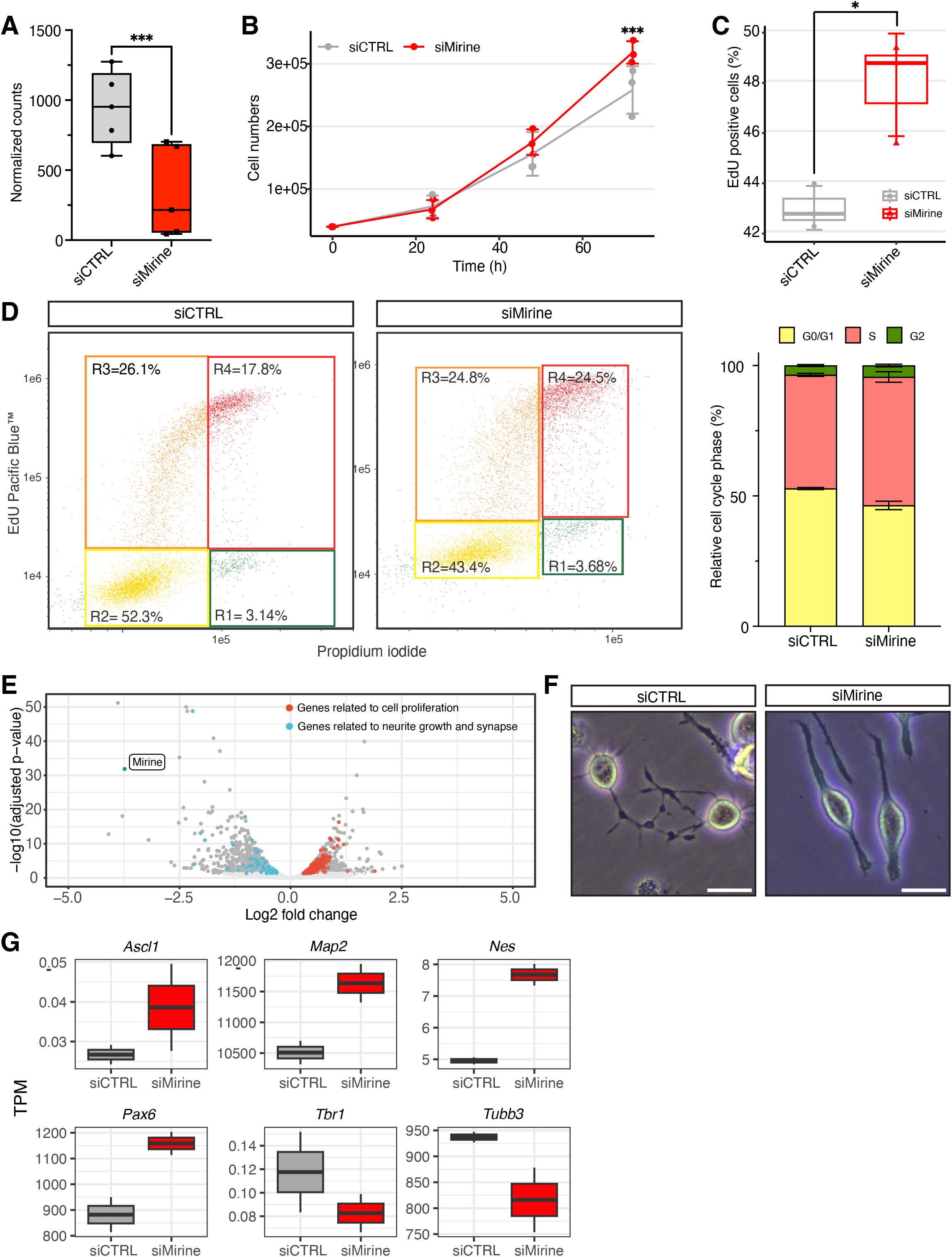
Function of MIRINE in balance between proliferation and differentiation. (A) Confirmation of *Mirine* mRNA knockdown using siRNA in N2A cells (n = 5; paired t-test). (B) Growth rates of *Mirine* knockdown cells (n = 3; two-way ANOVA). (C) Portion of proliferating cells (n = 3; unpaired t-test) and (D) cell cycle phase 48 h after *Mirine* knockdown (n = 3). (E) Differential gene expression of *Mirine* knockdown cells (n = 2). (F) Cellular morphology at 72 h after *Mirine* knockdown in N2A cells (n = 3; scale bar = 50 μm). (G) Representative expression of neuron-specific genes in si*Mirine*-transfected N2A cells (n = 2) (*p < 0.05; ***p < 0.001).

### *Mirine* knockdown leads to an impairment of neuronal differentiation *in vivo*

Based on the expression profile of *Mirine* in embryonic mouse brains (Figure 2B), we hypothesized that MIRINE may act during neurogenesis to induce or facilitate maturation of neuronal progenitors in the developing cerebral cortex. To test this, we performed *in-utero* electroporation (IUE) in E14.5 mouse cortices to express shRNA constructs targeting the CDS (sh*Mirine1*) or 3’UTR region (sh*Mirine2*) of *Mirine* within their developing cerebral cortex (Figure 4A). First, we wanted to establish that *in vivo* knockdown of *Mirine* recapitulates the proliferation phenotype we observed in cell culture. Therefore, we harvested E16.5 IUE mice forebrains 1 h after injecting EdU to visualize proliferating cells. Consistent with the N2A cells, sh*Mirine* cells (GFP^+^) showed twofold more EdU incorporation (GFP^+^EdU^+^) than the control in the fraction of IUE cortices (Figure 4B). Next, we focused on the differentiation status of sh*Mirine* neurons. We isolated primary cortical neurons from E15.5 mice, cultured them for 7-9 days (DIV) and performed a Sholl analysis to quantify the degree of neurite outgrowth. sh*Mirine* neurons showed significantly less complexity (Figure 4C), and a reduced number of neurites compared to control neurons (Figure 4D), indicating less complex dendritic branching. However, the length of axons was not affected by *Mirine* knockdown (Figure 4E). To examine the consequences of aberrant cortical neurogenesis upon knockdown of *Mirine in vivo*, we analyzed IUE mice two days after birth (P2) to assess the laminar position of targeted projection neurons. However, we did not observe a striking difference in the cohort (GFP^+^) of targeted projection neurons in the cortices of P2 mice, suggesting that their migration was not altered by a knockdown of *Mirine* (Supplementary Figure 5). This indicates that downregulation of *Mirine* leads to a defect of neurogenesis but does not delay neuronal migration in the cortex. In contrast, morpholino-induced knockdown of *mirine* in zebrafish embryos reduced the size of the forebrain significantly at day 5 post fertilization (dpf) while the midbrain or the eyes were not affected, indicating that a global reduction of *mirine* can affect brain development *in vivo* (Figure 4F). Together, our *in vivo* analyses showed that MIRINE promotes neuronal differentiation during neurogenesis and that its absence impacts forebrain development.

**Figure 4.**
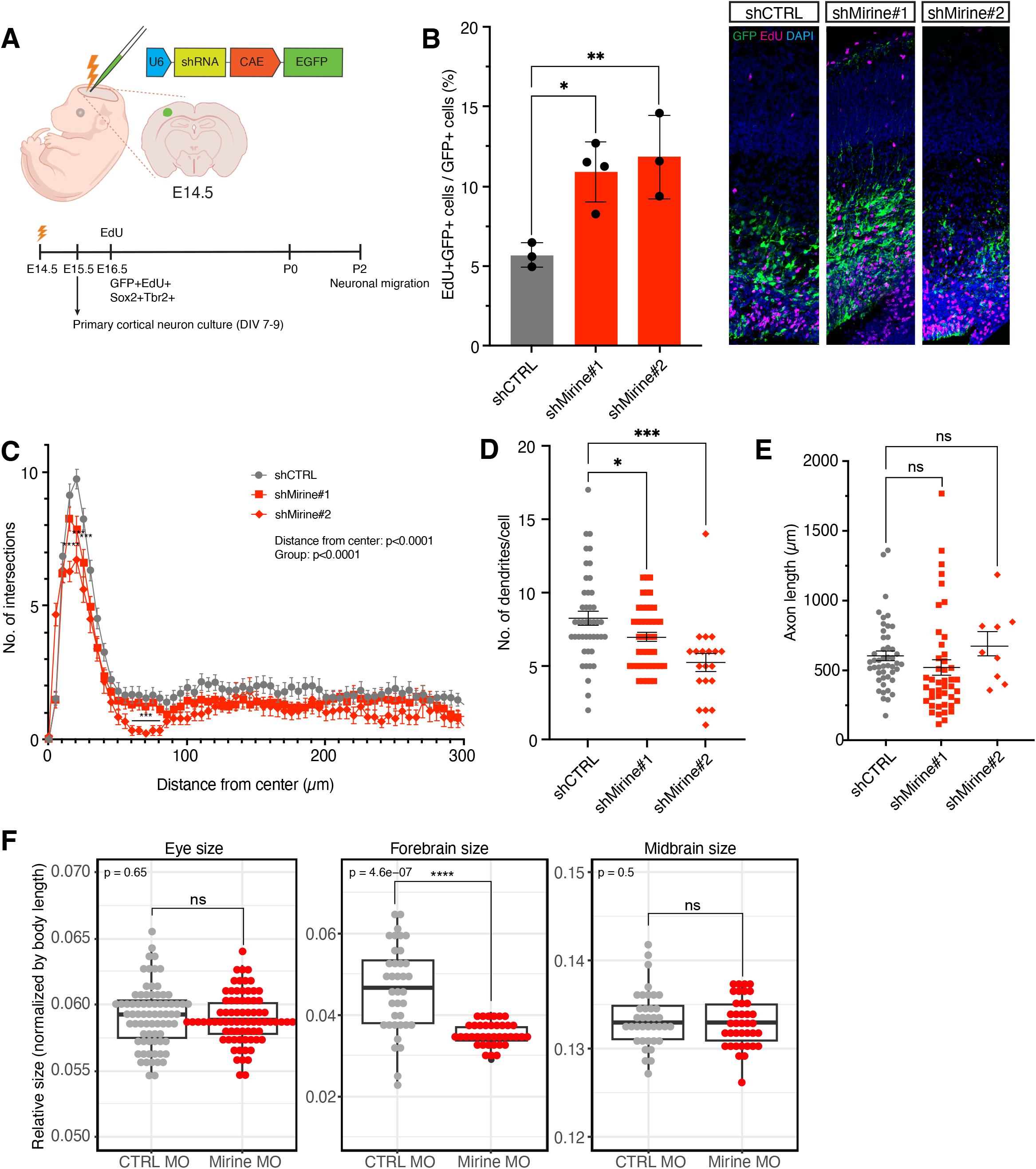
*In vivo* function of MIRINE in mouse neurogenesis. (A) Schematic representation of *in utero* knockdown experiment in E14.5 mouse embryonic cortices. (B) EdU incorporation in *Mirine* knockdown neurons at the coronal plane of E16.5 mouse embryonic cortices. Staining: right (knockdown cells (green); EdU (red); nuclei counterstained with DAPI (blue)). Double positive cells indicate proliferating cells. Quantification (left; n = 3-4; ANOVA with multiple comparisons). (C) Sholl analysis of DIV7-9 *Mirine* knockdown mouse primary neurons (n = 46 (control), 42 (shRNA#1), 8 (shRNA#2); two-way ANOVA). (D) Number of dendrites per cell and (E) axonal lengths in DIV 7-9 *Mirine* knockdown mouse primary neurons (n = 46 (control), 42 (shRNA#1), 8 (shRNA#2); ANOVA with multiple comparisons). (F) Eye size, forebrain size, and midbrain size after *mirine*-morpholino injection into zebrafish embryos (n=37; unpaired t-test) (*p < 0.05; **p < 0.01; *** p <0.001; ****p < 0.0001)

### MIRINE interacts with synaptic vesicle protein SYNGR3

To gain mechanistic insight into MIRINE function, we sought to identify its interacting partners. Our RNA-seq analysis of si*Mirine* N2A cells showed an enrichment of GO terms linked to protein transport and membrane-associated processes (Supplementary Figure 4A), suggesting a role of MIRINE in membrane protein regulation. To test whether MIRINE interacts with other membrane proteins, we performed an interactome analysis by expressing FLAG-tagged MIRINE in N2A cells, followed by immunoprecipitation and mass spectrometry (IP-MS). We identified 116 proteins enriched in the MIRINE interactome. Out of these, 12 proteins were annotated as membrane-associated proteins or transporter proteins (Table 4, Figure 5A), indicating an enrichment for membrane-related functions. We tested several candidates by co-immunoprecipitation (co-IP) followed by western blot and confirmed Synaptogyrin-3 (SYNGR3) as an interaction partner (Figure 5B). Furthermore, we predicted MIRINE-SYNGR3 complexes in mouse, human and zebrafish. AlphaFold predicted their binding via the C-terminal transmembrane domain of MIRINE with a high confidence (>70 pLDDT; Supplementary Figure 7A-C). The binding via the transmembrane domain is further supported by the finding that SYNGR3 is only detected in the co-IP of full length MIRINE but not when the transmembrane domain was truncated (Supplementary Figure 7D). Interestingly, SYNGR3 protein levels were reduced in si*Mirine* N2A cells, suggesting that MIRINE contributes to the stability or maintenance of SYNGR3 (Figure 5C). Furthermore, we observed a decrease in SYNGR3 levels in transfected primary neurons from mouse brain further supporting the functional connection (Figure 5D). In addition to *Syngr3*, several genes encoding proteins characteristic of synaptic vesicles such as *Sv2a*, *Sv2b*, *Syn1*, *Syp*, *Syt1*, *Syt7*, and *Vamp1* were downregulated in si*Mirine* cells (Figure 5E). Given the interaction between MIRINE and SYNGR3, we examined whether the downregulation of SYNGR3 phenocopies the knockdown of *Mirine* and performed siRNA-mediated *Syngr3* depletion in N2A cells. We observed that si*Syngr3* cells proliferate faster than control cells even though less pronounced than in *Mirine* knockdown (Figure 5F).

**Figure 5.**
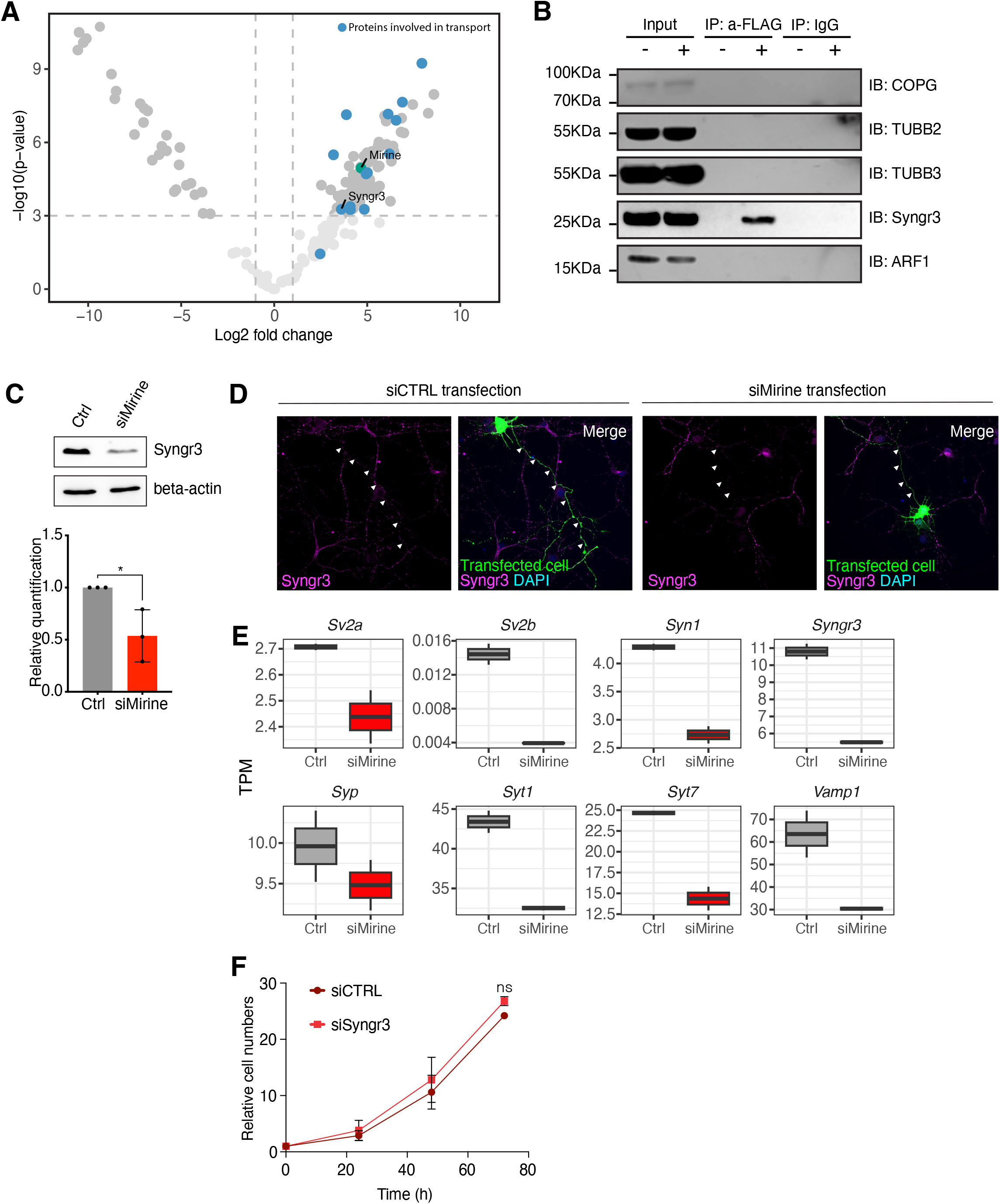
Interaction partner of MIRINE. (A) Interactome (IP-MS) of MIRINE in N2A cells; blue dots indicate proteins implicated in protein trafficking and membrane (n = 4). (B) Binding of MIRINE-interaction candidates by Co-IP using 3X-FLAG-MIRINE in N2A cells. (C) Expression of *Syngr3* in *Mirine* knockdown N2A cells (top) with quantification (bottom; n = 3; unpaired t-test) (D) Localization of SYNGR3 in DIV 7-9 *Mirine* knockdown mouse primary neurons. GFP^+^ indicates *Mirine* knockdown cells; arrowheads indicate axons. (E) mRNA expression of representative synaptic vesicle proteins in *Mirine* knockdown N2A cells (n = 2). (F) Cell growth analysis of *Syngr3* knockdown in N2A cells (n = 3) (ns, not significant; *p < 0.05).

**Table 4.**
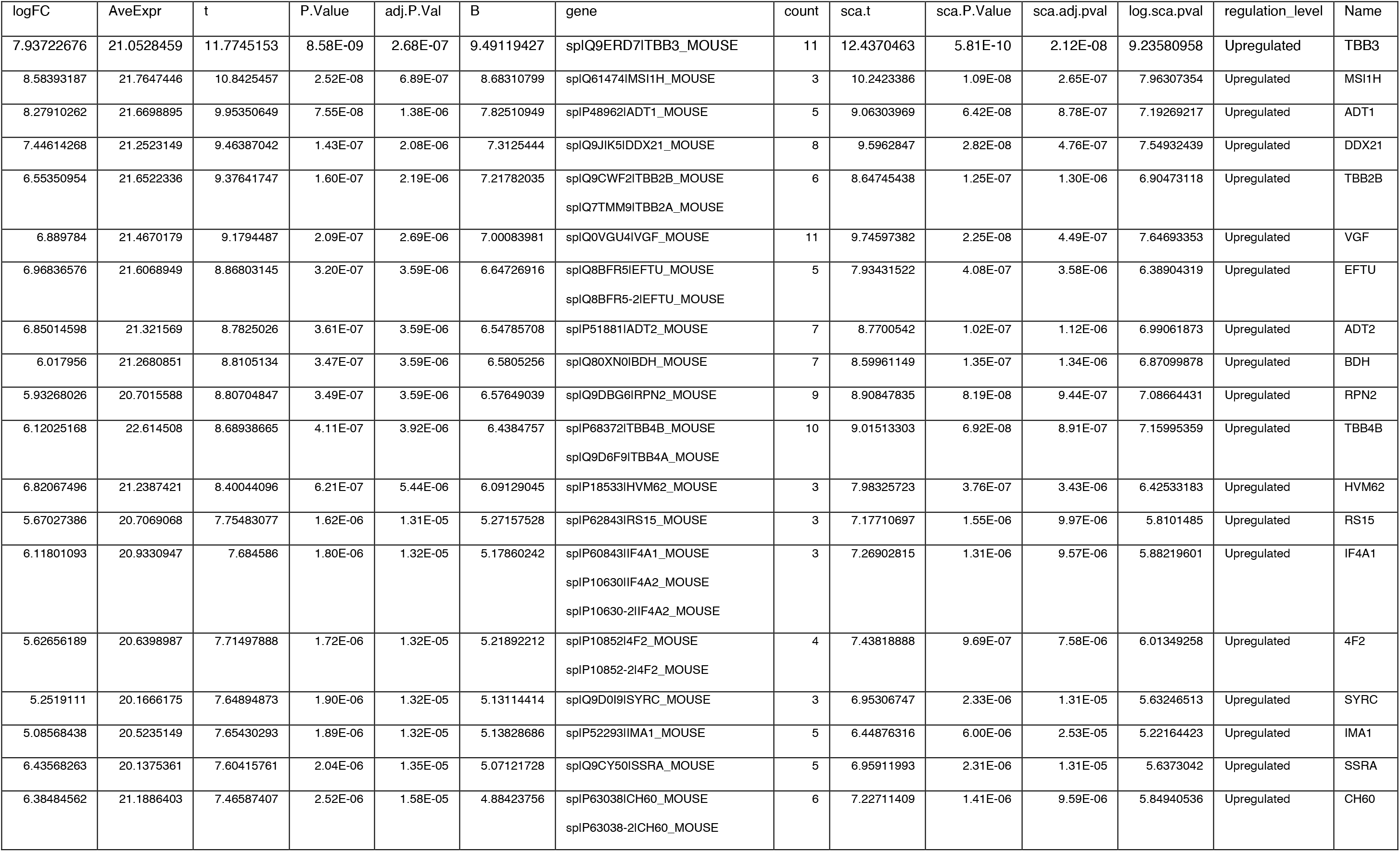
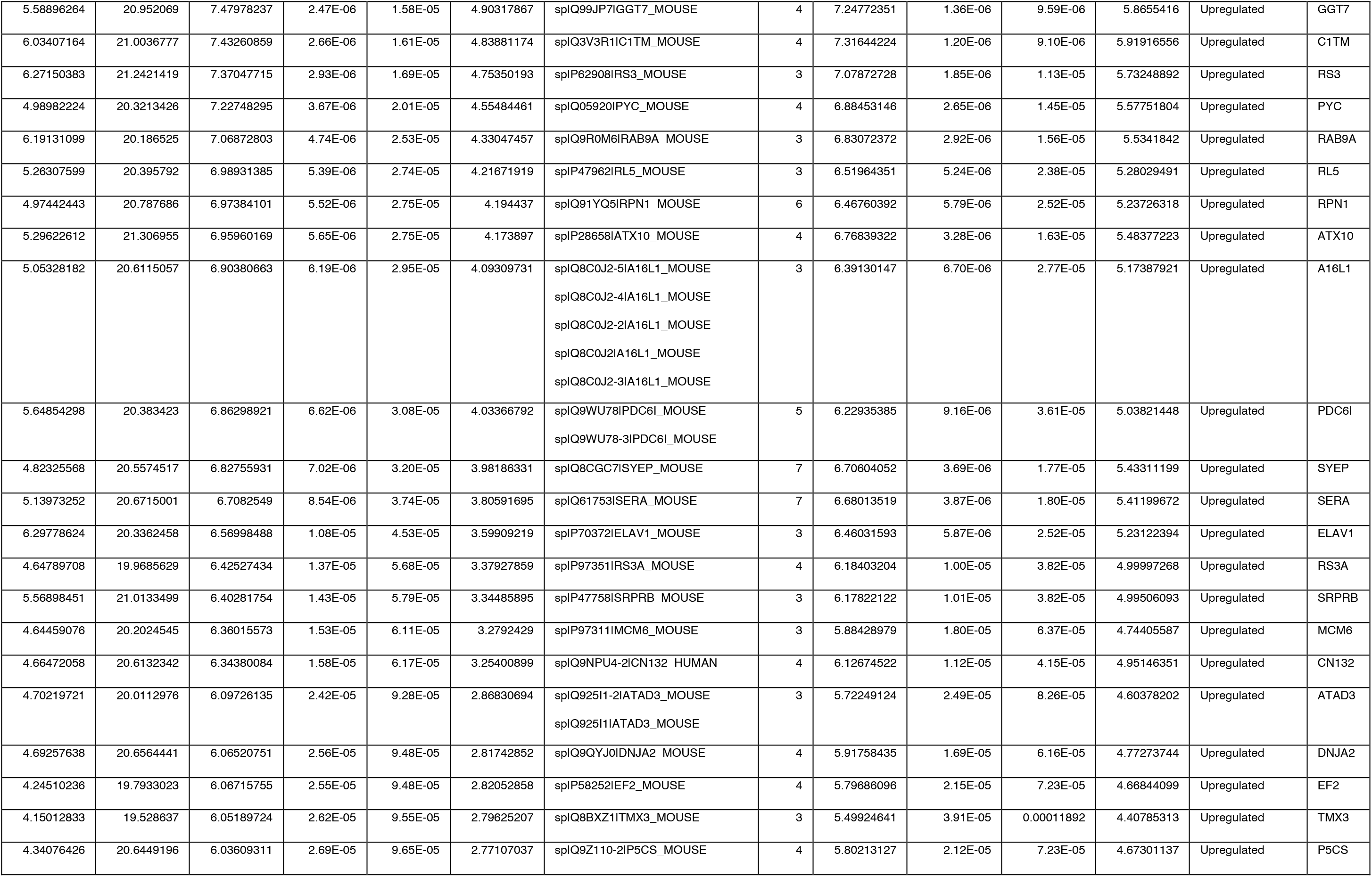
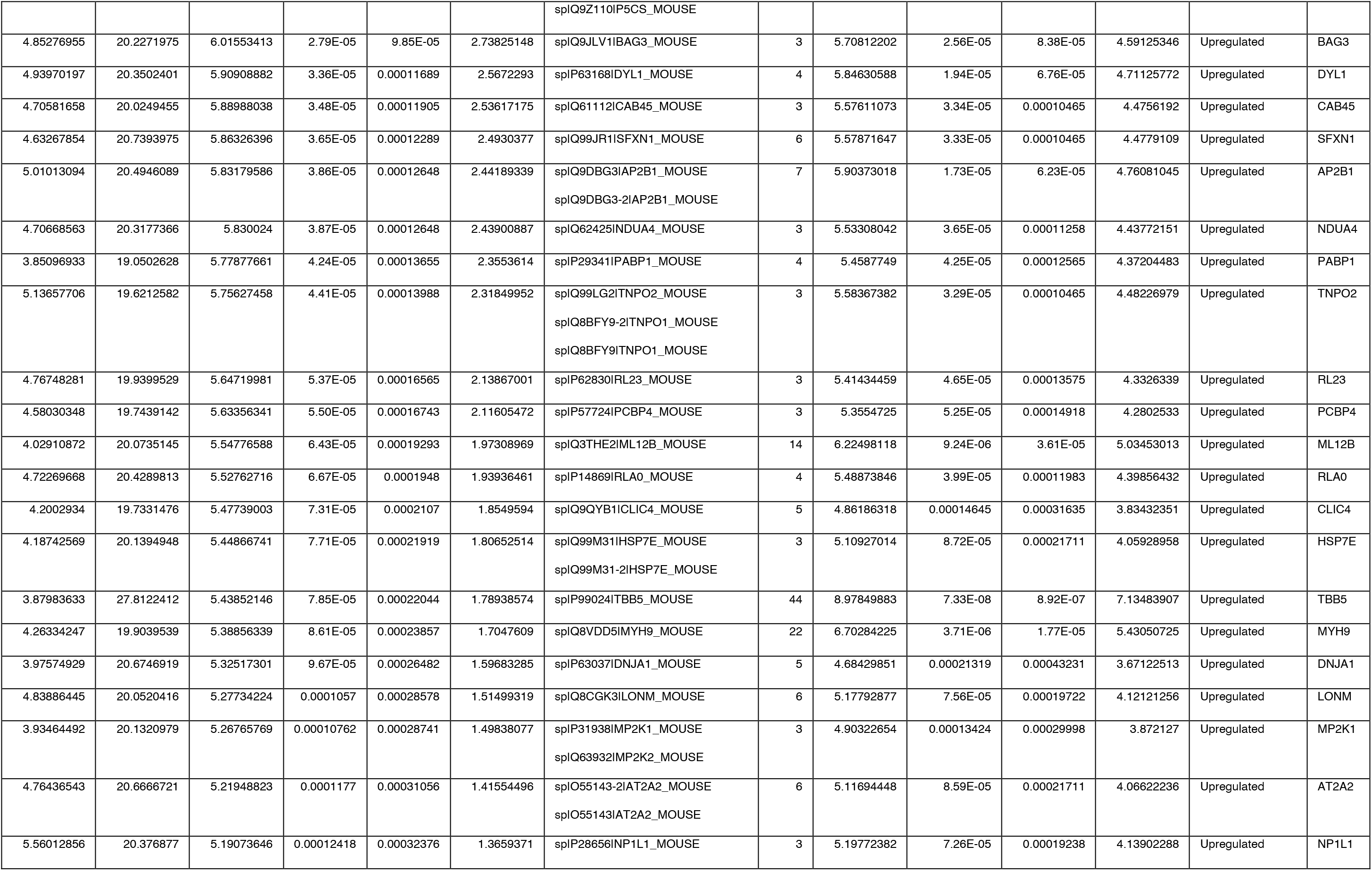
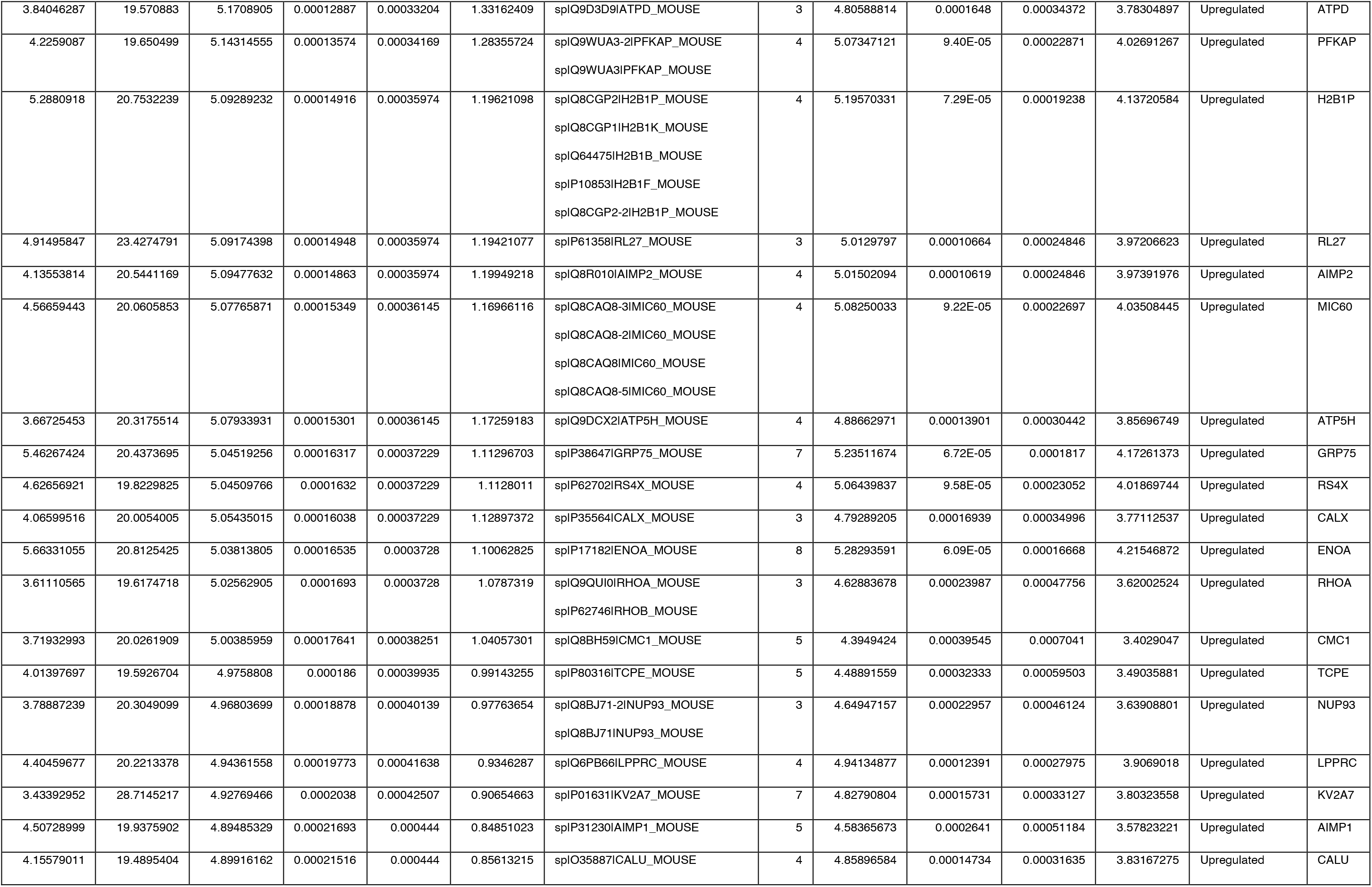
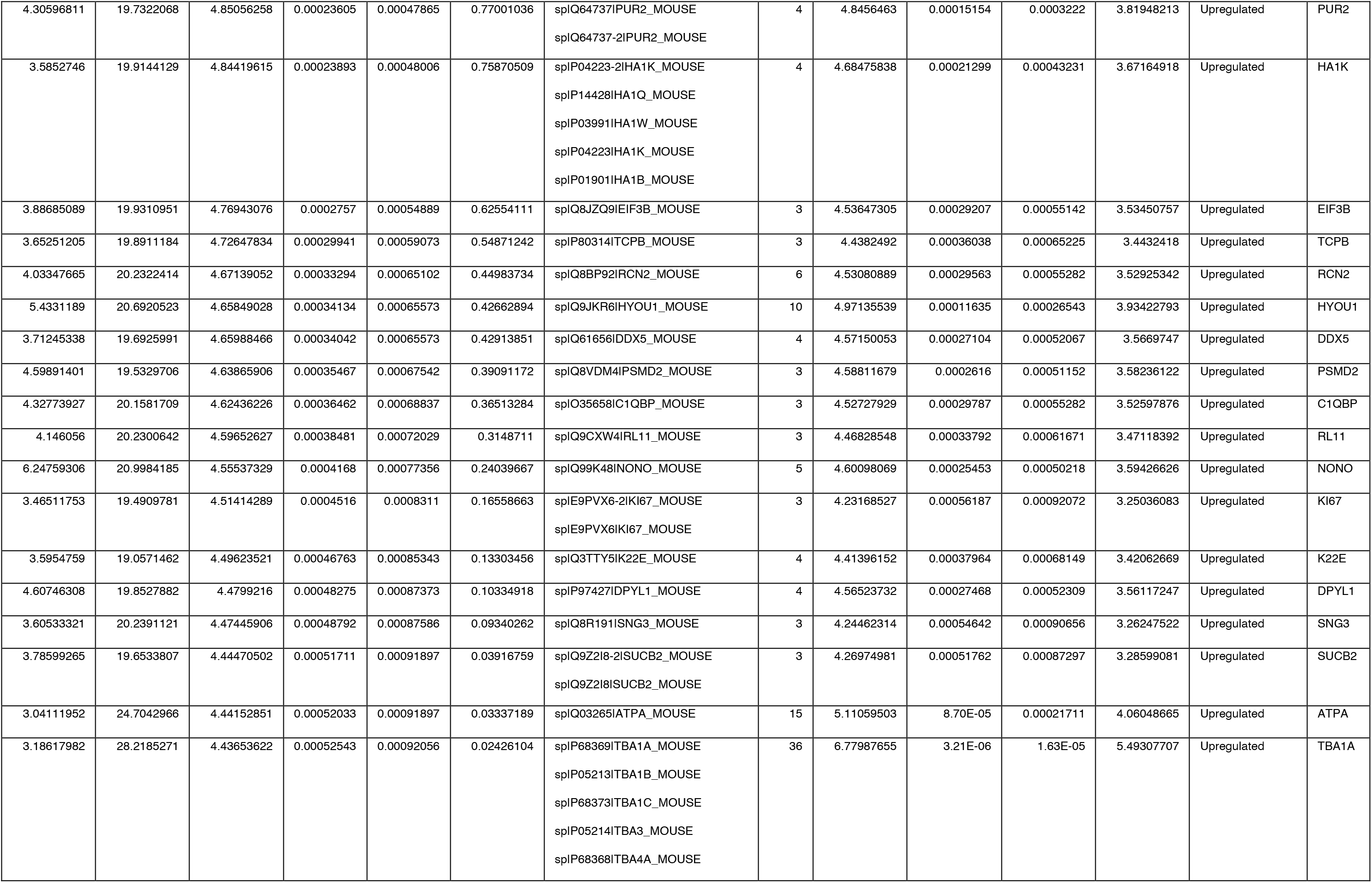
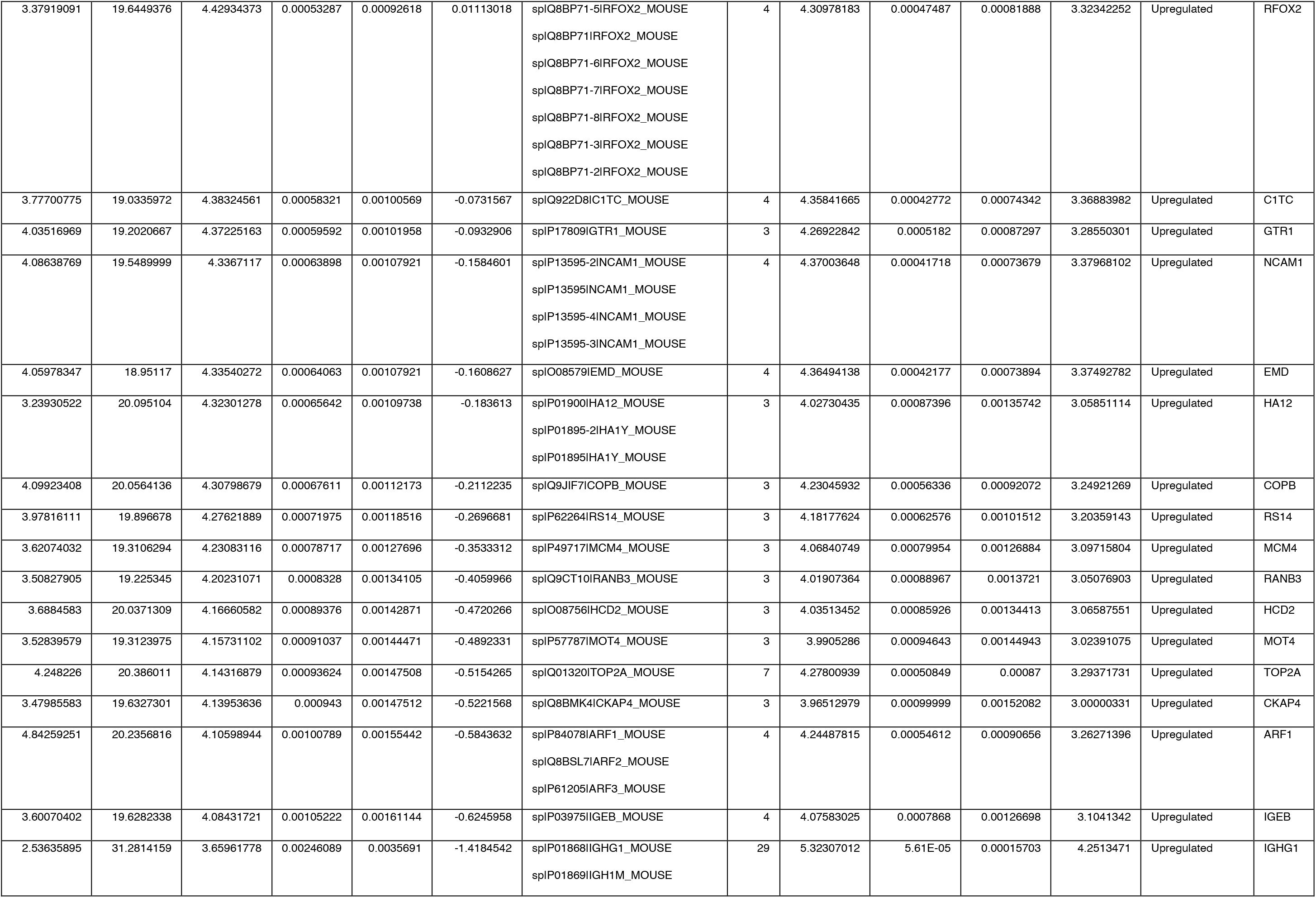
Enriched protein lists of IP-MS in N2A cells.

| logFC | AveExpr | t | P.Value | adj.P.Val | B | gene | count | sca.t | sca.P.Value | sca.adj.pval | log.sca.pval | regulation_level | Name |
|---|---|---|---|---|---|---|---|---|---|---|---|---|---|
| 7.93722676 | 21.0528459 | 11.7745153 | 8.58E-09 | 2.68E-07 | 9.49119427 | sp\|Q9ERD7\|TBB3_MOUSE | 11 | 12.4370463 | 5.81E-10 | 2.12E-08 | 9.23580958 | Upregulated | TBB3 |
| 8.58393187 | 21.7647446 | 10.8425457 | 2.52E-08 | 6.89E-07 | 8.68310799 | sp\|Q61474\|MSI1H_MOUSE | 3 | 10.2423386 | 1.09E-08 | 2.65E-07 | 7.96307354 | Upregulated | MSI1H |
| 8.27910262 | 21.6698895 | 9.95350649 | 7.55E-08 | 1.38E-06 | 7.82510949 | sp\|P48962\|ADT1_MOUSE | 5 | 9.06303969 | 6.42E-08 | 8.78E-07 | 7.19269217 | Upregulated | ADT1 |
| 7.44614268 | 21.2523149 | 9.46387042 | 1.43E-07 | 2.08E-06 | 7.3125444 | sp\|Q9JIK5\|DDX21_MOUSE | 8 | 9.5962847 | 2.82E-08 | 4.76E-07 | 7.54932439 | Upregulated | DDX21 |
| 6.55350954 | 21.6522336 | 9.37641747 | 1.60E-07 | 2.19E-06 | 7.21782035 | sp\|Q9CWF2\|TBB2B_MOUSE<br>sp\|Q7TMM9\|TBB2A_MOUSE | 6 | 8.64745438 | 1.25E-07 | 1.30E-06 | 6.90473118 | Upregulated | TBB2B |
| 6.889784 | 21.4670179 | 9.1794487 | 2.09E-07 | 2.69E-06 | 7.00083981 | sp\|Q0VGU4\|VGF_MOUSE | 11 | 9.74597382 | 2.25E-08 | 4.49E-07 | 7.64693353 | Upregulated | VGF |
| 6.96836576 | 21.6068949 | 8.86803145 | 3.20E-07 | 3.59E-06 | 6.64726916 | sp\|Q8BFR5\|EFTU_MOUSE<br>sp\|Q8BFR5-2\|EFTU_MOUSE | 5 | 7.93431522 | 4.08E-07 | 3.58E-06 | 6.38904319 | Upregulated | EFTU |
| 6.85014598 | 21.321569 | 8.7825026 | 3.61E-07 | 3.59E-06 | 6.54785708 | sp\|P51881\|ADT2_MOUSE | 7 | 8.7700542 | 1.02E-07 | 1.12E-06 | 6.99061873 | Upregulated | ADT2 |
| 6.017956 | 21.2680851 | 8.8105134 | 3.47E-07 | 3.59E-06 | 6.5805256 | sp\|Q80XN0\|BDH_MOUSE | 7 | 8.59961149 | 1.35E-07 | 1.34E-06 | 6.87099878 | Upregulated | BDH |
| 5.93268026 | 20.7015588 | 8.80704847 | 3.49E-07 | 3.59E-06 | 6.57649039 | sp\|Q9DBG6\|RPN2_MOUSE | 9 | 8.90847835 | 8.19E-08 | 9.44E-07 | 7.08664431 | Upregulated | RPN2 |
| 6.12025168 | 22.614508 | 8.68938665 | 4.11E-07 | 3.92E-06 | 6.4384757 | sp\|P68372\|TBB4B_MOUSE<br>sp\|Q9D6F9\|TBB4A_MOUSE | 10 | 9.01513303 | 6.92E-08 | 8.91E-07 | 7.15995359 | Upregulated | TBB4B |
| 6.82067496 | 21.2387421 | 8.40044096 | 6.21E-07 | 5.44E-06 | 6.09129045 | sp\|P18533\|HVM62_MOUSE | 3 | 7.98325723 | 3.76E-07 | 3.43E-06 | 6.42533183 | Upregulated | HVM62 |
| 5.67027386 | 20.7069068 | 7.75483077 | 1.62E-06 | 1.31E-05 | 5.27157528 | sp\|P62843\|RS15_MOUSE | 3 | 7.17710697 | 1.55E-06 | 9.97E-06 | 5.8101485 | Upregulated | RS15 |
| 6.11801093 | 20.9330947 | 7.684586 | 1.80E-06 | 1.32E-05 | 5.17860242 | sp\|P60843\|IF4A1_MOUSE<br>sp\|P10630\|IF4A2_MOUSE<br>sp\|P10630-2\|IF4A2_MOUSE | 3 | 7.26902815 | 1.31E-06 | 9.57E-06 | 5.88219601 | Upregulated | IF4A1 |
| 5.62656189 | 20.6398987 | 7.71497888 | 1.72E-06 | 1.32E-05 | 5.21892212 | sp\|P10852\|4F2_MOUSE<br>sp\|P10852-2\|4F2_MOUSE | 4 | 7.43818888 | 9.69E-07 | 7.58E-06 | 6.01349258 | Upregulated | 4F2 |
| 5.2519111 | 20.1666175 | 7.64894873 | 1.90E-06 | 1.32E-05 | 5.13114414 | sp\|Q9D0I9\|SYRC_MOUSE | 3 | 6.95306747 | 2.33E-06 | 1.31E-05 | 5.63246513 | Upregulated | SYRC |
| 5.08568438 | 20.5235149 | 7.65430293 | 1.89E-06 | 1.32E-05 | 5.13828686 | sp\|P52293\|IMA1_MOUSE | 5 | 6.44876316 | 6.00E-06 | 2.53E-05 | 5.22164423 | Upregulated | IMA1 |
| 6.43568263 | 20.1375361 | 7.60415761 | 2.04E-06 | 1.35E-05 | 5.07121728 | sp\|Q9CY50\|SSRA_MOUSE | 5 | 6.95911993 | 2.31E-06 | 1.31E-05 | 5.6373042 | Upregulated | SSRA |
| 6.38484562 | 21.1886403 | 7.46587407 | 2.52E-06 | 1.58E-05 | 4.88423756 | sp\|P63038\|CH60_MOUSE<br>sp\|P63038-2\|CH60_MOUSE | 6 | 7.22711409 | 1.41E-06 | 9.59E-06 | 5.84940536 | Upregulated | CH60 |
| 5.58896264 | 20.952069 | 7.47978237 | 2.47E-06 | 1.58E-05 | 4.90317867 | sp\|Q99JP7\|GGT7_MOUSE | 4 | 7.24772351 | 1.36E-06 | 9.59E-06 | 5.8655416 | Upregulated | GGT7 |
| 6.03407164 | 21.0036777 | 7.43260859 | 2.66E-06 | 1.61E-05 | 4.83881174 | sp\|Q3V3R1\|C1TM_MOUSE | 4 | 7.31644224 | 1.20E-06 | 9.10E-06 | 5.91916556 | Upregulated | C1TM |
| 6.27150383 | 21.2421419 | 7.37047715 | 2.93E-06 | 1.69E-05 | 4.75350193 | sp\|P62908\|RS3_MOUSE | 3 | 7.07872728 | 1.85E-06 | 1.13E-05 | 5.73248892 | Upregulated | RS3 |
| 4.98982224 | 20.3213426 | 7.22748295 | 3.67E-06 | 2.01E-05 | 4.55484461 | sp\|Q05920\|PYC_MOUSE | 4 | 6.88453146 | 2.65E-06 | 1.45E-05 | 5.57751804 | Upregulated | PYC |
| 6.19131099 | 20.186525 | 7.06872803 | 4.74E-06 | 2.53E-05 | 4.33047457 | sp\|Q9R0M6\|RAB9A_MOUSE | 3 | 6.83072372 | 2.92E-06 | 1.56E-05 | 5.5341842 | Upregulated | RAB9A |
| 5.26307599 | 20.395792 | 6.98931385 | 5.39E-06 | 2.74E-05 | 4.21671919 | sp\|P47962\|RL5_MOUSE | 3 | 6.51964351 | 5.24E-06 | 2.38E-05 | 5.28029491 | Upregulated | RL5 |
| 4.97442443 | 20.787686 | 6.97384101 | 5.52E-06 | 2.75E-05 | 4.194437 | sp\|Q91YQ5\|RPN1_MOUSE | 6 | 6.46760392 | 5.79E-06 | 2.52E-05 | 5.23726318 | Upregulated | RPN1 |
| 5.29622612 | 21.306955 | 6.95960169 | 5.65E-06 | 2.75E-05 | 4.173897 | sp\|P28658\|ATX10_MOUSE | 4 | 6.76839322 | 3.28E-06 | 1.63E-05 | 5.48377223 | Upregulated | ATX10 |
| 5.05328182 | 20.6115057 | 6.90380663 | 6.19E-06 | 2.95E-05 | 4.09309731 | sp\|Q8C0J2-5\|A16L1_MOUSE<br>sp\|Q8C0J2-4\|A16L1_MOUSE<br>sp\|Q8C0J2-2\|A16L1_MOUSE<br>sp\|Q8C0J2\|A16L1_MOUSE<br>sp\|Q8C0J2-3\|A16L1_MOUSE | 3 | 6.39130147 | 6.70E-06 | 2.77E-05 | 5.17387921 | Upregulated | A16L1 |
| 5.64854298 | 20.383423 | 6.86298921 | 6.62E-06 | 3.08E-05 | 4.03366792 | sp\|Q9WU78\|PDC6I_MOUSE<br>sp\|Q9WU78-3\|PDC6I_MOUSE | 5 | 6.22935385 | 9.16E-06 | 3.61E-05 | 5.03821448 | Upregulated | PDC6I |
| 4.82325568 | 20.5574517 | 6.82755931 | 7.02E-06 | 3.20E-05 | 3.98186331 | sp\|Q8CGC7\|SYEP_MOUSE | 7 | 6.70604052 | 3.69E-06 | 1.77E-05 | 5.43311199 | Upregulated | SYEP |
| 5.13973252 | 20.6715001 | 6.7082549 | 8.54E-06 | 3.74E-05 | 3.80591695 | sp\|Q61753\|SERA_MOUSE | 7 | 6.68013519 | 3.87E-06 | 1.80E-05 | 5.41199672 | Upregulated | SERA |
| 6.29778624 | 20.3362458 | 6.56998488 | 1.08E-05 | 4.53E-05 | 3.59909219 | sp\|P70372\|ELAV1_MOUSE | 3 | 6.46031593 | 5.87E-06 | 2.52E-05 | 5.23122394 | Upregulated | ELAV1 |
| 4.64789708 | 19.9685629 | 6.42527434 | 1.37E-05 | 5.68E-05 | 3.37927859 | sp\|P97351\|RS3A_MOUSE | 4 | 6.18403204 | 1.00E-05 | 3.82E-05 | 4.99997268 | Upregulated | RS3A |
| 5.56898451 | 21.0133499 | 6.40281754 | 1.43E-05 | 5.79E-05 | 3.34485895 | sp\|P47758\|SRPRB_MOUSE | 3 | 6.17822122 | 1.01E-05 | 3.82E-05 | 4.99506093 | Upregulated | SRPRB |
| 4.64459076 | 20.2024545 | 6.36015573 | 1.53E-05 | 6.11E-05 | 3.2792429 | sp\|P97311\|MCM6_MOUSE | 3 | 5.88428979 | 1.80E-05 | 6.37E-05 | 4.74405587 | Upregulated | MCM6 |
| 4.66472058 | 20.6132342 | 6.34380084 | 1.58E-05 | 6.17E-05 | 3.25400899 | sp\|Q9NPU4-2\|CN132_HUMAN | 4 | 6.12674522 | 1.12E-05 | 4.15E-05 | 4.95146351 | Upregulated | CN132 |
| 4.70219721 | 20.0112976 | 6.09726135 | 2.42E-05 | 9.28E-05 | 2.86830694 | sp\|Q925I1-2\|ATAD3_MOUSE<br>sp\|Q925I1\|ATAD3_MOUSE | 3 | 5.72249124 | 2.49E-05 | 8.26E-05 | 4.60378202 | Upregulated | ATAD3 |
| 4.69257638 | 20.6564441 | 6.06520751 | 2.56E-05 | 9.48E-05 | 2.81742852 | sp\|Q9QYJ0\|DNJA2_MOUSE | 4 | 5.91758435 | 1.69E-05 | 6.16E-05 | 4.77273744 | Upregulated | DNJA2 |
| 4.24510236 | 19.7933023 | 6.06715755 | 2.55E-05 | 9.48E-05 | 2.82052858 | sp\|P58252\|EF2_MOUSE | 4 | 5.79686096 | 2.15E-05 | 7.23E-05 | 4.66844099 | Upregulated | EF2 |
| 4.15012833 | 19.528637 | 6.05189724 | 2.62E-05 | 9.55E-05 | 2.79625207 | sp\|Q8BXZ1\|TMX3_MOUSE | 3 | 5.49924641 | 3.91E-05 | 0.00011892 | 4.40785313 | Upregulated | TMX3 |
| 4.34076426 | 20.6449196 | 6.03609311 | 2.69E-05 | 9.65E-05 | 2.77107037 | sp\|Q9Z110-2\|P5CS_MOUSE<br>sp\|Q9Z110\|P5CS_MOUSE | 4 | 5.80213127 | 2.12E-05 | 7.23E-05 | 4.67301137 | Upregulated | P5CS |
| 4.85276955 | 20.2271975 | 6.01553413 | 2.79E-05 | 9.85E-05 | 2.73825148 | sp\|Q9JLV1\|BAG3_MOUSE | 3 | 5.70812202 | 2.56E-05 | 8.38E-05 | 4.59125346 | Upregulated | BAG3 |
| 4.93970197 | 20.3502401 | 5.90908882 | 3.36E-05 | 0.00011689 | 2.5672293 | sp\|P63168\|DYL1_MOUSE | 4 | 5.84630588 | 1.94E-05 | 6.76E-05 | 4.71125772 | Upregulated | DYL1 |
| 4.70581658 | 20.0249455 | 5.88988038 | 3.48E-05 | 0.00011905 | 2.53617175 | sp\|Q61112\|CAB45_MOUSE | 3 | 5.57611073 | 3.34E-05 | 0.00010465 | 4.4756192 | Upregulated | CAB45 |
| 4.63267854 | 20.7393975 | 5.86326396 | 3.65E-05 | 0.00012289 | 2.4930377 | sp\|Q99JR1\|SFXN1_MOUSE | 6 | 5.57871647 | 3.33E-05 | 0.00010465 | 4.4779109 | Upregulated | SFXN1 |
| 5.01013094 | 20.4946089 | 5.83179586 | 3.86E-05 | 0.00012648 | 2.44189339 | sp\|Q9DBG3\|AP2B1_MOUSE<br>sp\|Q9DBG3-2\|AP2B1_MOUSE | 7 | 5.90373018 | 1.73E-05 | 6.23E-05 | 4.76081045 | Upregulated | AP2B1 |
| 4.70668563 | 20.3177366 | 5.830024 | 3.87E-05 | 0.00012648 | 2.43900887 | sp\|Q62425\|NDUA4_MOUSE | 3 | 5.53308042 | 3.65E-05 | 0.00011258 | 4.43772151 | Upregulated | NDUA4 |
| 3.85096933 | 19.0502628 | 5.77877661 | 4.24E-05 | 0.00013655 | 2.3553614 | sp\|P29341\|PABP1_MOUSE | 4 | 5.4587749 | 4.25E-05 | 0.00012565 | 4.37204483 | Upregulated | PABP1 |
| 5.13657706 | 19.6212582 | 5.75627458 | 4.41E-05 | 0.00013988 | 2.31849952 | sp\|Q99LG2\|TNPO2_MOUSE<br>sp\|Q8BFY9-2\|TNPO1_MOUSE<br>sp\|Q8BFY9\|TNPO1_MOUSE | 3 | 5.58367382 | 3.29E-05 | 0.00010465 | 4.48226979 | Upregulated | TNPO2 |
| 4.76748281 | 19.9399529 | 5.64719981 | 5.37E-05 | 0.00016565 | 2.13867001 | sp\|P62830\|RL23_MOUSE | 3 | 5.41434459 | 4.65E-05 | 0.00013575 | 4.3326339 | Upregulated | RL23 |
| 4.58030348 | 19.7439142 | 5.63356341 | 5.50E-05 | 0.00016743 | 2.11605472 | sp\|P57724\|PCBP4_MOUSE | 3 | 5.3554725 | 5.25E-05 | 0.00014918 | 4.2802533 | Upregulated | PCBP4 |
| 4.02910872 | 20.0735145 | 5.54776588 | 6.43E-05 | 0.00019293 | 1.97308969 | sp\|Q3THE2\|ML12B_MOUSE | 14 | 6.22498118 | 9.24E-06 | 3.61E-05 | 5.03453013 | Upregulated | ML12B |
| 4.72269668 | 20.4289813 | 5.52762716 | 6.67E-05 | 0.0001948 | 1.93936461 | sp\|P14869\|RLA0_MOUSE | 4 | 5.48873846 | 3.99E-05 | 0.00011983 | 4.39856432 | Upregulated | RLA0 |
| 4.2002934 | 19.7331476 | 5.47739003 | 7.31E-05 | 0.0002107 | 1.8549594 | sp\|Q9QYB1\|CLIC4_MOUSE | 5 | 4.86186318 | 0.00014645 | 0.00031635 | 3.83432351 | Upregulated | CLIC4 |
| 4.18742569 | 20.1394948 | 5.44866741 | 7.71E-05 | 0.00021919 | 1.80652514 | sp\|Q99M31\|HSP7E_MOUSE<br>sp\|Q99M31-2\|HSP7E_MOUSE | 3 | 5.10927014 | 8.72E-05 | 0.00021711 | 4.05928958 | Upregulated | HSP7E |
| 3.87983633 | 27.8122412 | 5.43852146 | 7.85E-05 | 0.00022044 | 1.78938574 | sp\|P99024\|TBB5_MOUSE | 44 | 8.97849883 | 7.33E-08 | 8.92E-07 | 7.13483907 | Upregulated | TBB5 |
| 4.26334247 | 19.9039539 | 5.38856339 | 8.61E-05 | 0.00023857 | 1.7047609 | sp\|Q8VDD5\|MYH9_MOUSE | 22 | 6.70284225 | 3.71E-06 | 1.77E-05 | 5.43050725 | Upregulated | MYH9 |
| 3.97574929 | 20.6746919 | 5.32517301 | 9.67E-05 | 0.00026482 | 1.59683285 | sp\|P63037\|DNJA1_MOUSE | 5 | 4.68429851 | 0.00021319 | 0.00043231 | 3.67122513 | Upregulated | DNJA1 |
| 4.83886445 | 20.0520416 | 5.27734224 | 0.0001057 | 0.00028578 | 1.51499319 | sp\|Q8CGK3\|LONM_MOUSE | 6 | 5.17792877 | 7.56E-05 | 0.00019722 | 4.12121256 | Upregulated | LONM |
| 3.93464492 | 20.1320979 | 5.26765769 | 0.00010762 | 0.00028741 | 1.49838077 | sp\|P31938\|MP2K1_MOUSE<br>sp\|Q63932\|MP2K2_MOUSE | 3 | 4.90322654 | 0.00013424 | 0.00029998 | 3.872127 | Upregulated | MP2K1 |
| 4.76436543 | 20.6666721 | 5.21948823 | 0.0001177 | 0.00031056 | 1.41554496 | sp\|O55143-2\|AT2A2_MOUSE<br>sp\|O55143\|AT2A2_MOUSE | 6 | 5.11694448 | 8.59E-05 | 0.00021711 | 4.06622236 | Upregulated | AT2A2 |
| 5.56012856 | 20.376877 | 5.19073646 | 0.00012418 | 0.00032376 | 1.3659371 | sp\|P28656\|NP1L1_MOUSE | 3 | 5.19772382 | 7.26E-05 | 0.00019238 | 4.13902288 | Upregulated | NP1L1 |
| 3.84046287 | 19.570883 | 5.1708905 | 0.00012887 | 0.00033204 | 1.33162409 | sp\|Q9D3D9\|ATPD_MOUSE | 3 | 4.80588814 | 0.0001648 | 0.00034372 | 3.78304897 | Upregulated | ATPD |
| 4.2259087 | 19.650499 | 5.14314555 | 0.00013574 | 0.00034169 | 1.28355724 | sp\|Q9WUA3-2\|PFKAP_MOUSE<br>sp\|Q9WUA3\|PFKAP_MOUSE | 4 | 5.07347121 | 9.40E-05 | 0.00022871 | 4.02691267 | Upregulated | PFKAP |
| 5.2880918 | 20.7532239 | 5.09289232 | 0.00014916 | 0.00035974 | 1.19621098 | sp\|Q8CGP2\|H2B1P_MOUSE<br>sp\|Q8CGP1\|H2B1K_MOUSE<br>sp\|Q64475\|H2B1B_MOUSE<br>sp\|P10853\|H2B1F_MOUSE<br>sp\|Q8CGP2-2\|H2B1P_MOUSE | 4 | 5.19570331 | 7.29E-05 | 0.00019238 | 4.13720584 | Upregulated | H2B1P |
| 4.91495847 | 23.4274791 | 5.09174398 | 0.00014948 | 0.00035974 | 1.19421077 | sp\|P61358\|RL27_MOUSE | 3 | 5.0129797 | 0.00010664 | 0.00024846 | 3.97206623 | Upregulated | RL27 |
| 4.13553814 | 20.5441169 | 5.09477632 | 0.00014863 | 0.00035974 | 1.19949218 | sp\|Q8R010\|AIMP2_MOUSE | 4 | 5.01502094 | 0.00010619 | 0.00024846 | 3.97391976 | Upregulated | AIMP2 |
| 4.56659443 | 20.0605853 | 5.07765871 | 0.00015349 | 0.00036145 | 1.16966116 | sp\|Q8CAQ8-3\|MIC60_MOUSE<br>sp\|Q8CAQ8-2\|MIC60_MOUSE<br>sp\|Q8CAQ8\|MIC60_MOUSE<br>sp\|Q8CAQ8-5\|MIC60_MOUSE | 4 | 5.08250033 | 9.22E-05 | 0.00022697 | 4.03508445 | Upregulated | MIC60 |
| 3.66725453 | 20.3175514 | 5.07933931 | 0.00015301 | 0.00036145 | 1.17259183 | sp\|Q9DCX2\|ATP5H_MOUSE | 4 | 4.88662971 | 0.00013901 | 0.00030442 | 3.85696749 | Upregulated | ATP5H |
| 5.46267424 | 20.4373695 | 5.04519256 | 0.00016317 | 0.00037229 | 1.11296703 | sp\|P38647\|GRP75_MOUSE | 7 | 5.23511674 | 6.72E-05 | 0.0001817 | 4.17261373 | Upregulated | GRP75 |
| 4.62656921 | 19.8229825 | 5.04509766 | 0.0001632 | 0.00037229 | 1.1128011 | sp\|P62702\|RS4X_MOUSE | 4 | 5.06439837 | 9.58E-05 | 0.00023052 | 4.01869744 | Upregulated | RS4X |
| 4.06599516 | 20.0054005 | 5.05435015 | 0.00016038 | 0.00037229 | 1.12897372 | sp\|P35564\|CALX_MOUSE | 3 | 4.79289205 | 0.00016939 | 0.00034996 | 3.77112537 | Upregulated | CALX |
| 5.66331055 | 20.8125425 | 5.03813805 | 0.00016535 | 0.0003728 | 1.10062825 | sp\|P17182\|ENOA_MOUSE | 8 | 5.28293591 | 6.09E-05 | 0.00016668 | 4.21546872 | Upregulated | ENOA |
| 3.61110565 | 19.6174718 | 5.02562905 | 0.0001693 | 0.0003728 | 1.0787319 | sp\|Q9QUI0\|RHOA_MOUSE<br>sp\|P62746\|RHOB_MOUSE | 3 | 4.62883678 | 0.00023987 | 0.00047756 | 3.62002524 | Upregulated | RHOA |
| 3.71932993 | 20.0261909 | 5.00385959 | 0.00017641 | 0.00038251 | 1.04057301 | sp\|Q8BH59\|CMC1_MOUSE | 5 | 4.3949424 | 0.00039545 | 0.0007041 | 3.4029047 | Upregulated | CMC1 |
| 4.01397697 | 19.5926704 | 4.9758808 | 0.000186 | 0.00039935 | 0.99143255 | sp\|P80316\|TCPE_MOUSE | 5 | 4.48891559 | 0.00032333 | 0.00059503 | 3.49035881 | Upregulated | TCPE |
| 3.78887239 | 20.3049099 | 4.96803699 | 0.00018878 | 0.00040139 | 0.97763654 | sp\|Q8BJ71-2\|NUP93_MOUSE<br>sp\|Q8BJ71\|NUP93_MOUSE | 3 | 4.64947157 | 0.00022957 | 0.00046124 | 3.63908801 | Upregulated | NUP93 |
| 4.40459677 | 20.2213378 | 4.94361558 | 0.00019773 | 0.00041638 | 0.9346287 | sp\|Q6PB66\|LPPRC_MOUSE | 4 | 4.94134877 | 0.00012391 | 0.00027975 | 3.9069018 | Upregulated | LPPRC |
| 3.43392952 | 28.7145217 | 4.92769466 | 0.0002038 | 0.00042507 | 0.90654663 | sp\|P01631\|KV2A7_MOUSE | 7 | 4.82790804 | 0.00015731 | 0.00033127 | 3.80323558 | Upregulated | KV2A7 |
| 4.50728999 | 19.9375902 | 4.89485329 | 0.00021693 | 0.000444 | 0.84851023 | sp\|P31230\|AIMP1_MOUSE | 5 | 4.58365673 | 0.0002641 | 0.00051184 | 3.57823221 | Upregulated | AIMP1 |
| 4.15579011 | 19.4895404 | 4.89916162 | 0.00021516 | 0.000444 | 0.85613215 | sp\|O35887\|CALU_MOUSE | 4 | 4.85896584 | 0.00014734 | 0.00031635 | 3.83167275 | Upregulated | CALU |
| 4.30596811 | 19.7322068 | 4.85056258 | 0.00023605 | 0.00047865 | 0.77001036 | sp\|Q64737\|PUR2_MOUSE<br>sp\|Q64737-2\|PUR2_MOUSE | 4 | 4.8456463 | 0.00015154 | 0.0003222 | 3.81948213 | Upregulated | PUR2 |
| 3.5852746 | 19.9144129 | 4.84419615 | 0.00023893 | 0.00048006 | 0.75870509 | sp\|P04223-2\|HA1K_MOUSE<br>sp\|P14428\|HA1Q_MOUSE<br>sp\|P03991\|HA1W_MOUSE<br>sp\|P04223\|HA1K_MOUSE<br>sp\|P01901\|HA1B_MOUSE | 4 | 4.68475838 | 0.00021299 | 0.00043231 | 3.67164918 | Upregulated | HA1K |
| 3.88685089 | 19.9310951 | 4.76943076 | 0.0002757 | 0.00054889 | 0.62554111 | sp\|Q8JZQ9\|EIF3B_MOUSE | 3 | 4.53647305 | 0.00029207 | 0.00055142 | 3.53450757 | Upregulated | EIF3B |
| 3.65251205 | 19.8911184 | 4.72647834 | 0.00029941 | 0.00059073 | 0.54871242 | sp\|P80314\|TCPB_MOUSE | 3 | 4.4382492 | 0.00036038 | 0.00065225 | 3.4432418 | Upregulated | TCPB |
| 4.03347665 | 20.2322414 | 4.67139052 | 0.00033294 | 0.00065102 | 0.44983734 | sp\|Q8BP92\|RCN2_MOUSE | 6 | 4.53080889 | 0.00029563 | 0.00055282 | 3.52925342 | Upregulated | RCN2 |
| 5.4331189 | 20.6920523 | 4.65849028 | 0.00034134 | 0.00065573 | 0.42662894 | sp\|Q9JKR6\|HYOU1_MOUSE | 10 | 4.97135539 | 0.00011635 | 0.00026543 | 3.93422793 | Upregulated | HYOU1 |
| 3.71245338 | 19.6925991 | 4.65988466 | 0.00034042 | 0.00065573 | 0.42913851 | sp\|Q61656\|DDX5_MOUSE | 4 | 4.57150053 | 0.00027104 | 0.00052067 | 3.5669747 | Upregulated | DDX5 |
| 4.59891401 | 19.5329706 | 4.63865906 | 0.00035467 | 0.00067542 | 0.39091172 | sp\|Q8VDM4\|PSMD2_MOUSE | 3 | 4.58811679 | 0.0002616 | 0.00051152 | 3.58236122 | Upregulated | PSMD2 |
| 4.32773927 | 20.1581709 | 4.62436226 | 0.00036462 | 0.00068837 | 0.36513284 | sp\|O35658\|C1QBP_MOUSE | 3 | 4.52727929 | 0.00029787 | 0.00055282 | 3.52597876 | Upregulated | C1QBP |
| 4.146056 | 20.2300642 | 4.59652627 | 0.00038481 | 0.00072029 | 0.3148711 | sp\|Q9CXW4\|RL11_MOUSE | 3 | 4.46828548 | 0.00033792 | 0.00061671 | 3.47118392 | Upregulated | RL11 |
| 6.24759306 | 20.9984185 | 4.55537329 | 0.0004168 | 0.00077356 | 0.24039667 | sp\|Q99K48\|NONO_MOUSE | 5 | 4.60098069 | 0.00025453 | 0.00050218 | 3.59426626 | Upregulated | NONO |
| 3.46511753 | 19.4909781 | 4.51414289 | 0.0004516 | 0.0008311 | 0.16558663 | sp\|E9PVX6-2\|KI67_MOUSE<br>sp\|E9PVX6\|KI67_MOUSE | 3 | 4.23168527 | 0.00056187 | 0.00092072 | 3.25036083 | Upregulated | KI67 |
| 3.5954759 | 19.0571462 | 4.49623521 | 0.00046763 | 0.00085343 | 0.13303456 | sp\|Q3TTY5\|K22E_MOUSE | 4 | 4.41396152 | 0.00037964 | 0.00068149 | 3.42062669 | Upregulated | K22E |
| 4.60746308 | 19.8527882 | 4.4799216 | 0.00048275 | 0.00087373 | 0.10334918 | sp\|P97427\|DPYL1_MOUSE | 4 | 4.56523732 | 0.00027468 | 0.00052309 | 3.56117247 | Upregulated | DPYL1 |
| 3.60533321 | 20.2391121 | 4.47445906 | 0.00048792 | 0.00087586 | 0.09340262 | sp\|Q8R191\|SNG3_MOUSE | 3 | 4.24462314 | 0.00054642 | 0.00090656 | 3.26247522 | Upregulated | SNG3 |
| 3.78599265 | 19.6533807 | 4.44470502 | 0.00051711 | 0.00091897 | 0.03916759 | sp\|Q9Z2I8-2\|SUCB2_MOUSE<br>sp\|Q9Z2I8\|SUCB2_MOUSE | 3 | 4.26974981 | 0.00051762 | 0.00087297 | 3.28599081 | Upregulated | SUCB2 |
| 3.04111952 | 24.7042966 | 4.44152851 | 0.00052033 | 0.00091897 | 0.03337189 | sp\|Q03265\|ATPA_MOUSE | 15 | 5.11059503 | 8.70E-05 | 0.00021711 | 4.06048665 | Upregulated | ATPA |
| 3.18617982 | 28.2185271 | 4.43653622 | 0.00052543 | 0.00092056 | 0.02426104 | sp\|P68369\|TBA1A_MOUSE<br>sp\|P05213\|TBA1B_MOUSE<br>sp\|P68373\|TBA1C_MOUSE<br>sp\|P05214\|TBA3_MOUSE<br>sp\|P68368\|TBA4A_MOUSE | 36 | 6.77987655 | 3.21E-06 | 1.63E-05 | 5.49307707 | Upregulated | TBA1A |
| 3.37919091 | 19.6449376 | 4.42934373 | 0.00053287 | 0.00092618 | 0.01113018 | sp\|Q8BP71-5\|RFOX2_MOUSE<br>sp\|Q8BP71\|RFOX2_MOUSE<br>sp\|Q8BP71-6\|RFOX2_MOUSE<br>sp\|Q8BP71-7\|RFOX2_MOUSE<br>sp\|Q8BP71-8\|RFOX2_MOUSE<br>sp\|Q8BP71-3\|RFOX2_MOUSE<br>sp\|Q8BP71-2\|RFOX2_MOUSE | 4 | 4.30978183 | 0.00047487 | 0.00081888 | 3.32342252 | Upregulated | RFOX2 |
| 3.77700775 | 19.0335972 | 4.38324561 | 0.00058321 | 0.00100569 | -0.0731567 | sp\|Q922D8\|C1TC_MOUSE | 4 | 4.35841665 | 0.00042772 | 0.00074342 | 3.36883982 | Upregulated | C1TC |
| 4.03516969 | 19.2020667 | 4.37225163 | 0.00059592 | 0.00101958 | -0.0932906 | sp\|P17809\|GTR1_MOUSE | 3 | 4.26922842 | 0.0005182 | 0.00087297 | 3.28550301 | Upregulated | GTR1 |
| 4.08638769 | 19.5489999 | 4.3367117 | 0.00063898 | 0.00107921 | -0.1584601 | sp\|P13595-2\|NCAM1_MOUSE<br>sp\|P13595\|NCAM1_MOUSE<br>sp\|P13595-4\|NCAM1_MOUSE<br>sp\|P13595-3\|NCAM1_MOUSE | 4 | 4.37003648 | 0.00041718 | 0.00073679 | 3.37968102 | Upregulated | NCAM1 |
| 4.05978347 | 18.95117 | 4.33540272 | 0.00064063 | 0.00107921 | -0.1608627 | sp\|O08579\|EMD_MOUSE | 4 | 4.36494138 | 0.00042177 | 0.00073894 | 3.37492782 | Upregulated | EMD |
| 3.23930522 | 20.095104 | 4.32301278 | 0.00065642 | 0.00109738 | -0.183613 | sp\|P01900\|HA12_MOUSE<br>sp\|P01895-2\|HA1Y_MOUSE<br>sp\|P01895\|HA1Y_MOUSE | 3 | 4.02730435 | 0.00087396 | 0.00135742 | 3.05851114 | Upregulated | HA12 |
| 4.09923408 | 20.0564136 | 4.30798679 | 0.00067611 | 0.00112173 | -0.2112235 | sp\|Q9JIF7\|COPB_MOUSE | 3 | 4.23045932 | 0.00056336 | 0.00092072 | 3.24921269 | Upregulated | COPB |
| 3.97816111 | 19.896678 | 4.27621889 | 0.00071975 | 0.00118516 | -0.2696681 | sp\|P62264\|RS14_MOUSE | 3 | 4.18177624 | 0.00062576 | 0.00101512 | 3.20359143 | Upregulated | RS14 |
| 3.62074032 | 19.3106294 | 4.23083116 | 0.00078717 | 0.00127696 | -0.3533312 | sp\|P49717\|MCM4_MOUSE | 3 | 4.06840749 | 0.00079954 | 0.00126884 | 3.09715804 | Upregulated | MCM4 |
| 3.50827905 | 19.225345 | 4.20231071 | 0.0008328 | 0.00134105 | -0.4059966 | sp\|Q9CT10\|RANB3_MOUSE | 3 | 4.01907364 | 0.00088967 | 0.0013721 | 3.05076903 | Upregulated | RANB3 |
| 3.6884583 | 20.0371309 | 4.16660582 | 0.00089376 | 0.00142871 | -0.4720266 | sp\|O08756\|HCD2_MOUSE | 3 | 4.03513452 | 0.00085926 | 0.00134413 | 3.06587551 | Upregulated | HCD2 |
| 3.52839579 | 19.3123975 | 4.15731102 | 0.00091037 | 0.00144471 | -0.4892331 | sp\|P57787\|MOT4_MOUSE | 3 | 3.9905286 | 0.00094643 | 0.00144943 | 3.02391075 | Upregulated | MOT4 |
| 4.248226 | 20.386011 | 4.14316879 | 0.00093624 | 0.00147508 | -0.5154265 | sp\|Q01320\|TOP2A_MOUSE | 7 | 4.27800939 | 0.00050849 | 0.00087 | 3.29371731 | Upregulated | TOP2A |
| 3.47985583 | 19.6327301 | 4.13953636 | 0.000943 | 0.00147512 | -0.5221568 | sp\|Q8BMK4\|CKAP4_MOUSE | 3 | 3.96512979 | 0.00099999 | 0.00152082 | 3.00000331 | Upregulated | CKAP4 |
| 4.84259251 | 20.2356816 | 4.10598944 | 0.00100789 | 0.00155442 | -0.5843632 | sp\|P84078\|ARF1_MOUSE<br>sp\|Q8BSL7\|ARF2_MOUSE<br>sp\|P61205\|ARF3_MOUSE | 4 | 4.24487815 | 0.00054612 | 0.00090656 | 3.26271396 | Upregulated | ARF1 |
| 3.60070402 | 19.6282338 | 4.08431721 | 0.00105222 | 0.00161144 | -0.6245958 | sp\|P03975\|IGEB_MOUSE | 4 | 4.07583025 | 0.0007868 | 0.00126698 | 3.1041342 | Upregulated | IGEB |
| 2.53635895 | 31.2814159 | 3.65961778 | 0.00246089 | 0.0035691 | -1.4184542 | sp\|P01868\|IGHG1_MOUSE<br>sp\|P01869\|IGH1M_MOUSE | 29 | 5.32307012 | 5.61E-05 | 0.00015703 | 4.2513471 | Upregulated | IGHG1 |

SYNGR3 interacts with MIRINE, its protein levels are decreased in si*Mirine* cells, and its absence phenocopies the growth phenotype of *Mirine* depletion. However, our interactome analysis identified 116 MIRINE-interacting proteins enriched for membrane proteins and transporters, suggesting that MIRINE regulates neuronal differentiation through stabilization of multiple membrane proteins rather than a single target.

## Discussion

We performed ribosome profiling and identified 198 and 121 smORF candidates in zebrafish and mouse brains, respectively. Based on conservation and functional predictions, we selected MIRINE, a conserved and brain-enriched SEP encoded by the smORF *C14orf132/D430019H16Rik*. The conservation of MIRINE across 205 vertebrate species strongly indicates its functional importance in vertebrates (Figure 1B). Unlike many previously identified SEPs that show limited phylogenetic conservation, MIRINE exhibits high sequence conservation, suggesting that this short protein fulfils an evolutionarily preserved function. Its brain-enriched expression pattern (Figures 1E and 2A) and enrichment in cortical and hippocampal neurons (Figure 2D) further support a role of MIRINE in neuronal development.

Functionally, we demonstrated that MIRINE acts as a regulator of the proliferation-versus-differentiation homeostasis during neuronal development. Knockdown of *Mirine* consistently enhanced proliferation in N2A cells and in the embryonic cortex *in vivo*, as shown by increased EdU incorporation and an altered cell cycle distribution (Figure 3C-D and Figure 4B), while the expression of neuronal differentiation markers and neurite complexity was reduced (Figure 3E-F). Single-cell RNA-seq analysis suggests that *Mirine* expression is low in radial glia, increased in neuroblasts and highest in neurons, supporting a role at the stage of neuronal commitment rather than in progenitor maintenance (Figure 2B). Consistent with this directionality, overexpression of *mirine* did not cause an obvious morphological phenotype in HuC-GFP zebrafish embryos (Supplementary Figure 6), suggesting that endogenous *mirine* levels are not rate-limiting for its physiological function, which may restrain proliferation rather than actively instruct differentiation.

Mechanistically, MIRINE likely functions through membrane-associated protein complexes. Its C-terminal transmembrane domain is essential for its membrane localization, as truncation of this domain abolishes membrane association (Figure 1D). Interactome analysis identified 116 candidate interactors, many of which are membrane proteins and transporters (Figure 5A and Table 4). Among these, we validated SYNGR3 as an interaction partner (Figure 5B). SYNGR3 protein levels were significantly reduced upon *Mirine* knockdown both in N2A cells and in mouse brain (Figure 5C-D), and siRNA-mediated depletion of *SYNGR3* partially phenocopied the proliferation effect of *Mirine* knockdown (Figure 5F), supporting a functional link between the two proteins. Moreover, downregulation of additional synaptic vesicle-associated transcripts (*Sv2a*, *Sv2b*, *Syn1*, *Syp*, *Syt1*, *Syt7*, *Vamp1*) in *Mirine* knockdown cells (Figure 5E) suggests that this functional mechanism is not restricted to SYNGR3 alone. MIRINE may interact indirectly with a broader group of membrane and vesicle-associated proteins rather than functioning through a single downstream target, although direct biochemical evidence is currently limited to SYNGR3, and the remaining interactome candidates have not been experimentally validated.

Our *in vivo* experiments in mouse and zebrafish further support a conserved physiological role of MIRINE. *In utero* electroporation of shRNA constructs targeting *Mirine* in mouse embryos recapitulated increased proliferation and impaired neuronal maturation *in vitro* (Figure 4C-F), and morpholino-mediated knockdown in zebrafish embryos reduced forebrain size without affecting the midbrain or eyes (Figure 4F), showing that MIRINE plays a conserved role in regulating the transition between proliferation and differentiation during vertebrate brain development.

While this study was in progress, SEP18 was reported to be encoded at the same gene locus. *Sep18* is expressed in vascular smooth muscle cells, where its knockout promotes cell proliferation and exacerbates carotid artery neointima formation (Li et al. 2023). Our observation of increasing proliferating cells upon *Mirine* depletion in neuronal cells *in vitro* and *in vivo* is consistent with their finding that *Sep18* knockout elevates EdU incorporation in vascular smooth muscle, indicating that MIRINE affects the regulation of cell proliferation, which is recapitulated by two independent systems (Li et al. 2023). However, the physiological context, function, and mechanism we report in this study are markedly different. While their study mainly focused on the function of the microprotein in the cardiovascular system, we focused on the dominant expression pattern of *Mirine* in the brain. These complementary findings suggest that a single conserved SEP can restrain proliferation across multiple tissues while engaging distinct effector pathways in a context-dependent manner. Interestingly, a *Sep18* knockout mouse exists (Li et al. 2023). It will therefore be very interesting to analyze the brain structure and in particular the cortical layers in these mutants.

### Limitation of Study

Several limitations of this study should be noted. First, most of our functional experiments relied on the functional knockdown of *Mirine* rather than the knockout. Likewise, *in utero* electroporation targets a limited cortical region. Therefore, we did not observe a universal knockdown of *Mirine* in the mouse brain to cross-validate the effect observed in zebrafish embryos upon morpholino knockdown. Second, although our interactome data provided 116 candidate interactors, validation of direct interaction was limited. Validating these candidates would further test our model that MIRINE acts through a broader set of membrane proteins.

Together, our findings identify MIRINE as a conserved membrane-associated SEP that controls neuronal lineage progression by balancing proliferation and differentiation, likely through coordinated stabilization of synaptic vesicle-associated membrane proteins, including its interacting partner, SYNGR3. Therefore, MIRINE extends the field of functional SEPs, by introducing a differentiation regulator of the nervous system.

## Acknowledgements

We thank Erez Raz for support and advice at an early stage of this project. We thank Nadia Mercader-Huber, Xavi Langa and Inês Marques for support and discussions, and all members of the Leidel lab for critical discussions and feedback. We thank Claudia Gräf for sequencing the ribosome profiling libraries and the Next Generation Sequencing Platform of the University of Bern for the excellent service in performing the high-throughput sequencing and the Core Facility Proteomics & Mass Spectrometry of the University of Bern for performing our proteomics experiments. Work in the Leidel laboratory was supported by the Canton of Bern. Y.K. was supported by Seal of Excellence Funding of the University of Bern (SELF2019-02) and the Postdoctoral Fellowship Program (Nurturing Next-generation Researchers) by the National Research Foundation of Korea (2020R1A6A3A03038387). Y.K. was part of the COMET program of the University of Bern. The Nguyen laboratory is supported by ULiège (Crédit Classique), the F.R.S.-FNRS (PDR T.0185.20; EOS 0019118F-RG36), the WEL Research Institute (CR-2022A-12), the Fonds Leon Fredericq, the Fondation Simone et Pierre Clerdent, the Fondation Médicale Reine Elisabeth, the ERANET Neuron (STEM-MCD and NeuroTalk), the Win2Wal (ChipOmics; #2010126), and the ERC-Synergy (UNFOLD). The graphical abstract and Figure 4A was created in BioRender.

## Data Availability

Deep-sequencing data are available at the Gene Expression Omnibus (GEO), accession number: GSE347702. Proteomics data are available via ProteomeXchange with identifier PXD086624.

## Author contribution

Y.K., conceptualization, methodology, investigation, formal analysis, writing – original draft, and funding acquisition; B.S.N., conceptualization, methodology, software, investigation, formal analysis, data curation; A.E., investigation, formal analysis; N.K., investigation, formal analysis; R.K., methodology, formal analysis, data curation; H.C.A.D., investigation, formal analysis, data curation; L.N., supervision, writing – review & editing, and funding acquisition; S.A.L., conceptualization, supervision, writing – review & editing, and funding acquisition

## Conflict-of-interest declarations

All authors declare no conflict of interests.

**Supplementary Figure 1. Identification of small ORFs in zebrafish (left) and mouse (right) ribosome profiling.** (A) Codon periodicity and (B) Read lengths of zebrafish and mouse ribosome profiling reads results. (C) Number of identified small ORFs based on their category. (D) ORF lengths of detected small ORFs. (E) Number of overlapped small ORFs among ribosome profiling, shotgun proteomics, phylogenetic conservation analysis, and computational domain prediction.

**Supplementary Figure 2. Characterization of MIRINE.** (A) *Mirine*/*mirine* expression level in human, mouse, and zebrafish organs. Data is obtained from dbGaP Accession phs000424.v8.p2. (B) Alpha-fold prediction of MIRINE protein structure. (C) Time-course *Mirine* RNA expression levels of developing mouse organs. Call sets are from the ENCODE portal (Sloan et al. 2016) (https://www.encodeproject.org/) with the following identifiers: ENCSR304RDL, ENCSR160IIN, ENCSR647QBV, ENCSR970EWM, ENCSR185LWM, ENCSR752RGN, ENCSR080EVZ, ENCSR362AIZ, ENCSR049UJU, ENCSR727FHP, ENCSR691OPQ, ENCSR526SEX, ENCSR020DGG, ENCSR597UZW, ENCSR284YKY, ENCSR150CUE, ENCSR000BYS, ENCSR000BZI, ENCSR648YEP, ENCSR096STK, ENCSR000BZH, ENCSR284AMY, ENCSR867YNV, ENCSR826HIQ, ENCSR000BZO, ENCSR448MXQ, ENCSR611PTP, ENCSR764OPZ, ENCSR307BCA, ENCSR367ZPZ, ENCSR908JWT, ENCSR719NAJ, ENCSR557RMA, ENCSR792RJV, ENCSR343YLB, ENCSR559TRB, ENCSR760TOE, ENCSR401BSG, ENCSR017JEG, ENCSR285WZV, ENCSR420QTO, ENCSR921PRX, ENCSR943LKA, ENCSR982MRY, ENCSR457RRW, ENCSR039ADS, ENCSR992WBR, ENCSR906YQZ, ENCSR290RRR, ENCSR178GUS, ENCSR466KZY, ENCSR173PJN, ENCSR537GNQ, ENCSR062VTB, ENCSR504GEG, ENCSR370SFB, ENCSR932TRU, ENCSR848GST, ENCSR331XCE.

**Supplementary Figure 3. Mirine expression in mouse brain.** (A) Developmental time-course expression of *Mirine* in diverse cell types in embryonic mouse brain. (B) *Mirine* expression levels in neuronal cells. (C) *Mirine* expression levels in glutamatergic and GABAergic neurons. (D) *In situ* hybridization of *Mirine* in adult mouse brain. Arrowheads indicate hippocampus and arrows indicate cortex. From: Allen Mouse Brain Atlas: https://mouse.brain-map.org/experiment/show/70795509. Statistical analysis: Mann-Whitney U test with Benjamini-Hochberg post procedure (B, C).

**Supplementary Figure 4. Characterization of *Mirine* knockdown N2A cells.** (A) Gene ontology analysis of *Mirine* knockdown N2A cells. (B) *Mirine* RNA expression in RA-treated differentiated N2A cells.

**Supplementary Figure 5. Cortical migration of sh*Mirine* neurons in mouse brain at P2.** Staining: knockdown cells (green); nuclei counterstained with DAPI (blue).

**Supplementary Figure 6. mirine overexpression in HuC-GFP zebrafish embryos.** Zebrafish embryos after injection of *mirine* mRNA at 24 hpf (A) and 48 hpf (B); brightfield (top); GFP (bottom). HuC expression indicates neurons.

**Supplementary Figure 7. Comparison of structural predictions of MIRINE-SYNGR3 complexes in selected organisms.** (A) MIRINE-SYNGR3 complexes from *Danio rerio* (left), *Homo sapiens* (middle) and *Mus musculus* (right) exhibit similar binding modes and relative orientations of N- and C-termini. Green, SYNGR3; orange, MIRINE. (B) Confidence of AlphaFold predictions shown in (A). Structural models are coloured according to Predicted Local Distance Difference Test (pLDDT) scores using the colour scale on the right. (C) Predicted Aligned Error (PAE) scores of corresponding models highlight similar MIRINE-SYNGR3 interaction mode via C-terminal transmembrane helix of MIRINE. (D) Detection of SYNGR3 in different co-IP experiments; label-free quantification (LFQ) values for each replicate (Full: full-length MIRINE; Trunc: transmembrane domain-truncated MIRINE; FLAG: co-IP using anti-FLAG antibody; IgG: co-IP using control IgG). Note that SYNGR3 is only detected if full length MIRINE is used as bait.

