## Supplementary Figures for "The small ORF (smORF)-encoded polypeptide MIRINE promotes neuronal differentiation in vertebrate brain development"

### Supplementary Figure 1

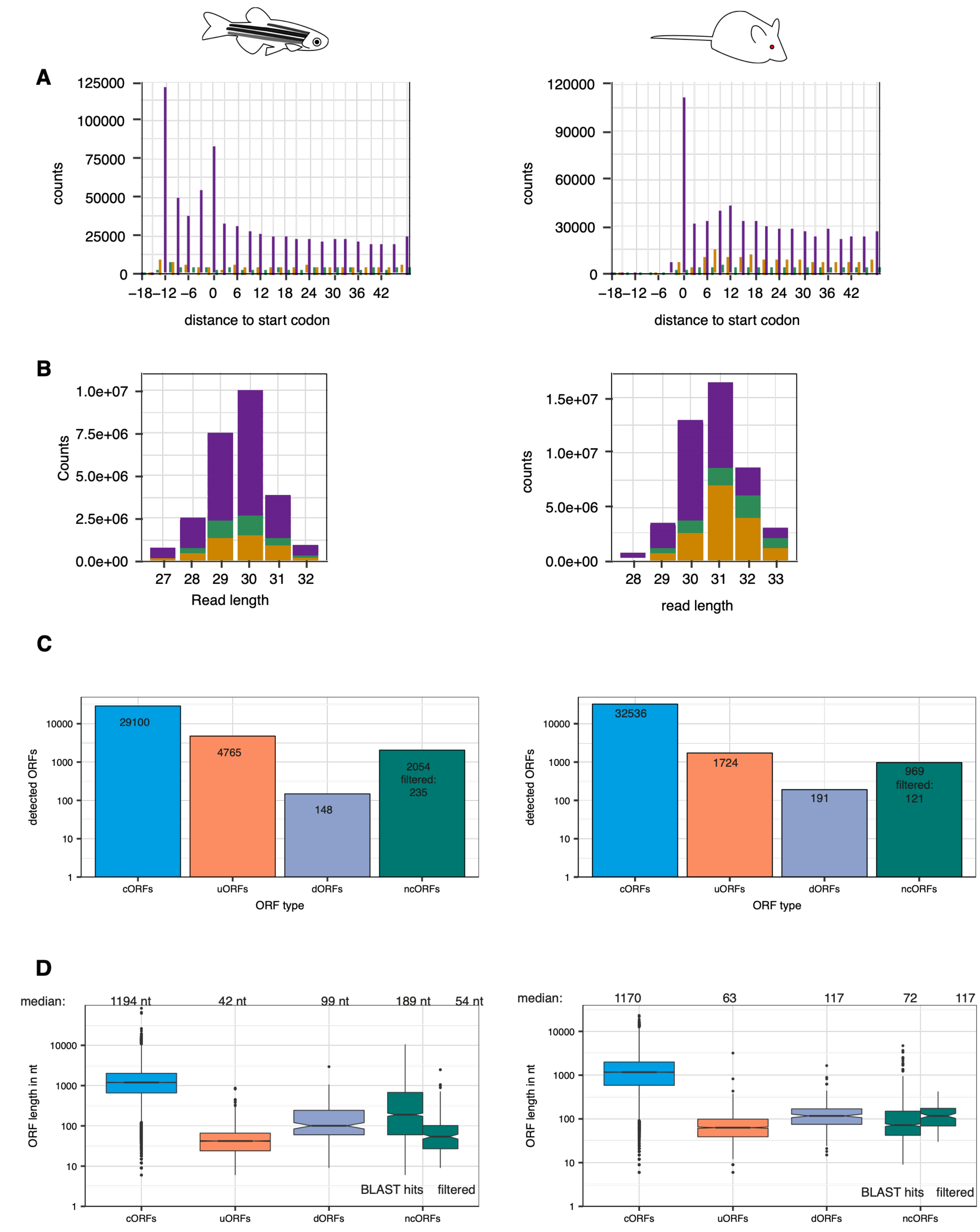

E

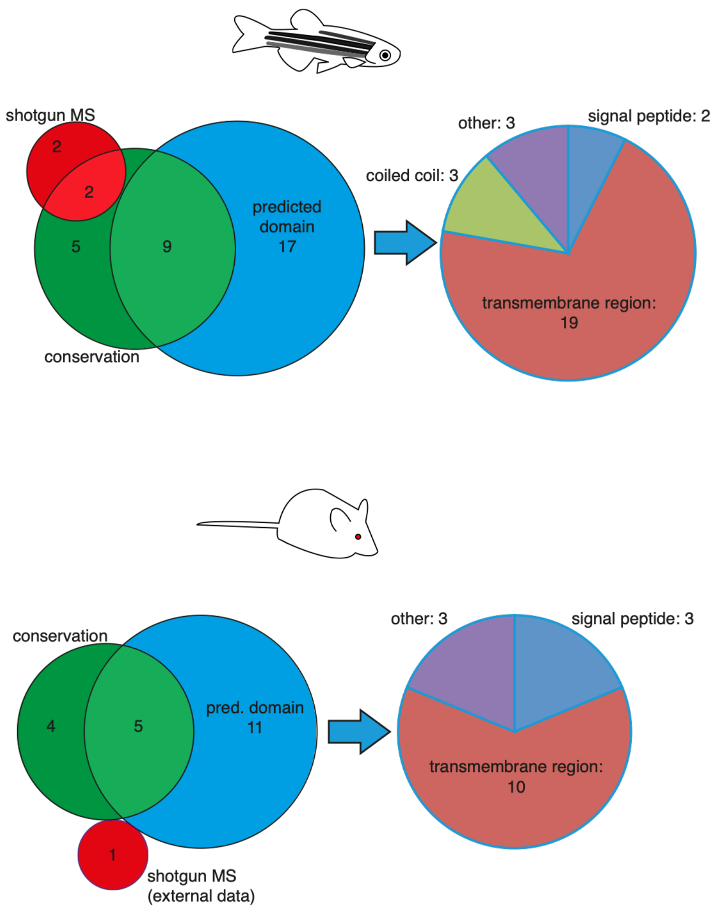
shotgun MS
2
2
5
9
predicted
domain
17
conservation
other: 3
signal peptide: 2
coiled coil: 3
transmembrane region:
19
conservation
4
5
pred. domain
11
1
shotgun MS
(external data)
other: 3
signal peptide: 3
transmembrane region:
10

#### Supplementary Figure 2

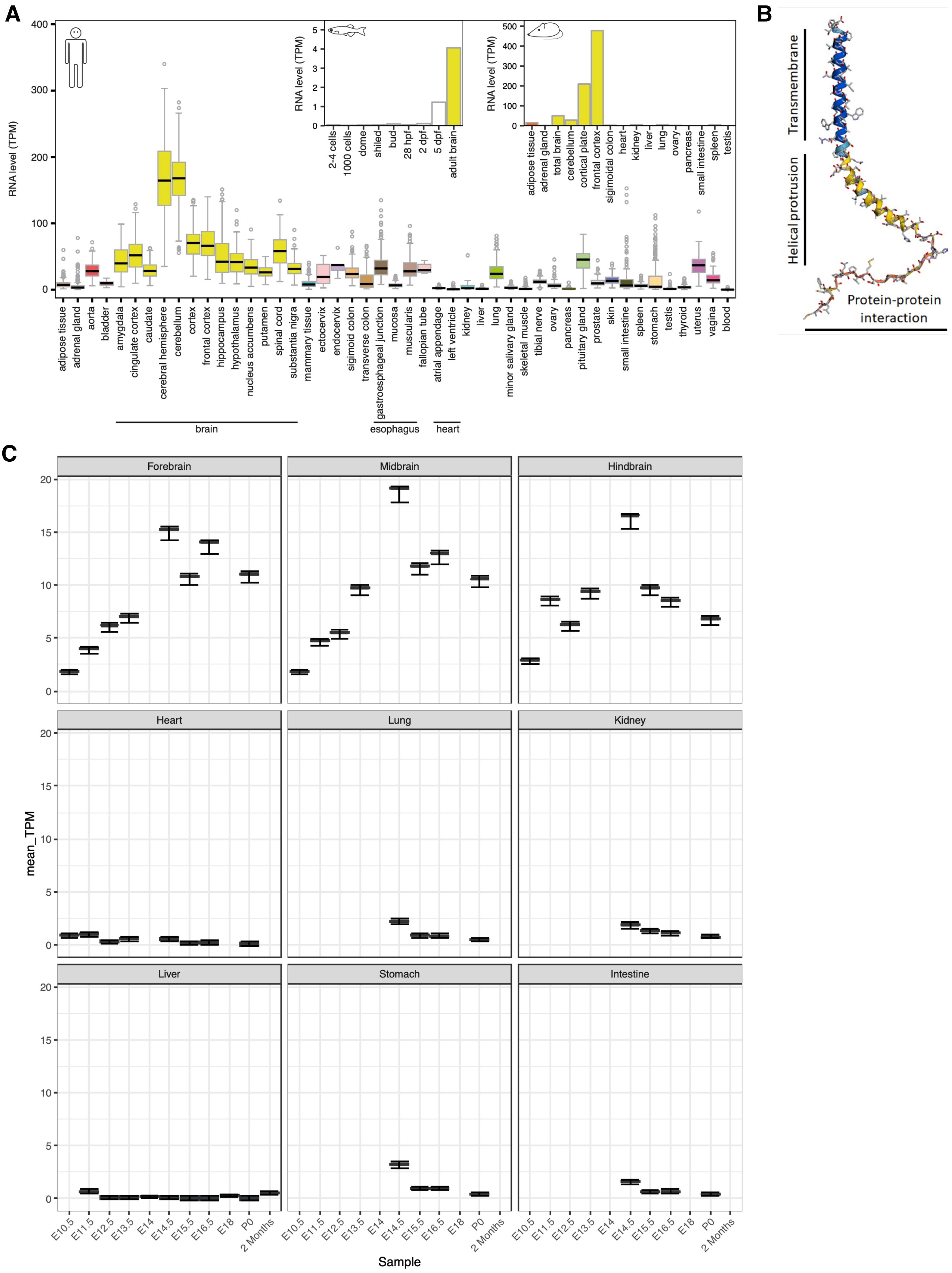

#### Supplementary Figure 3

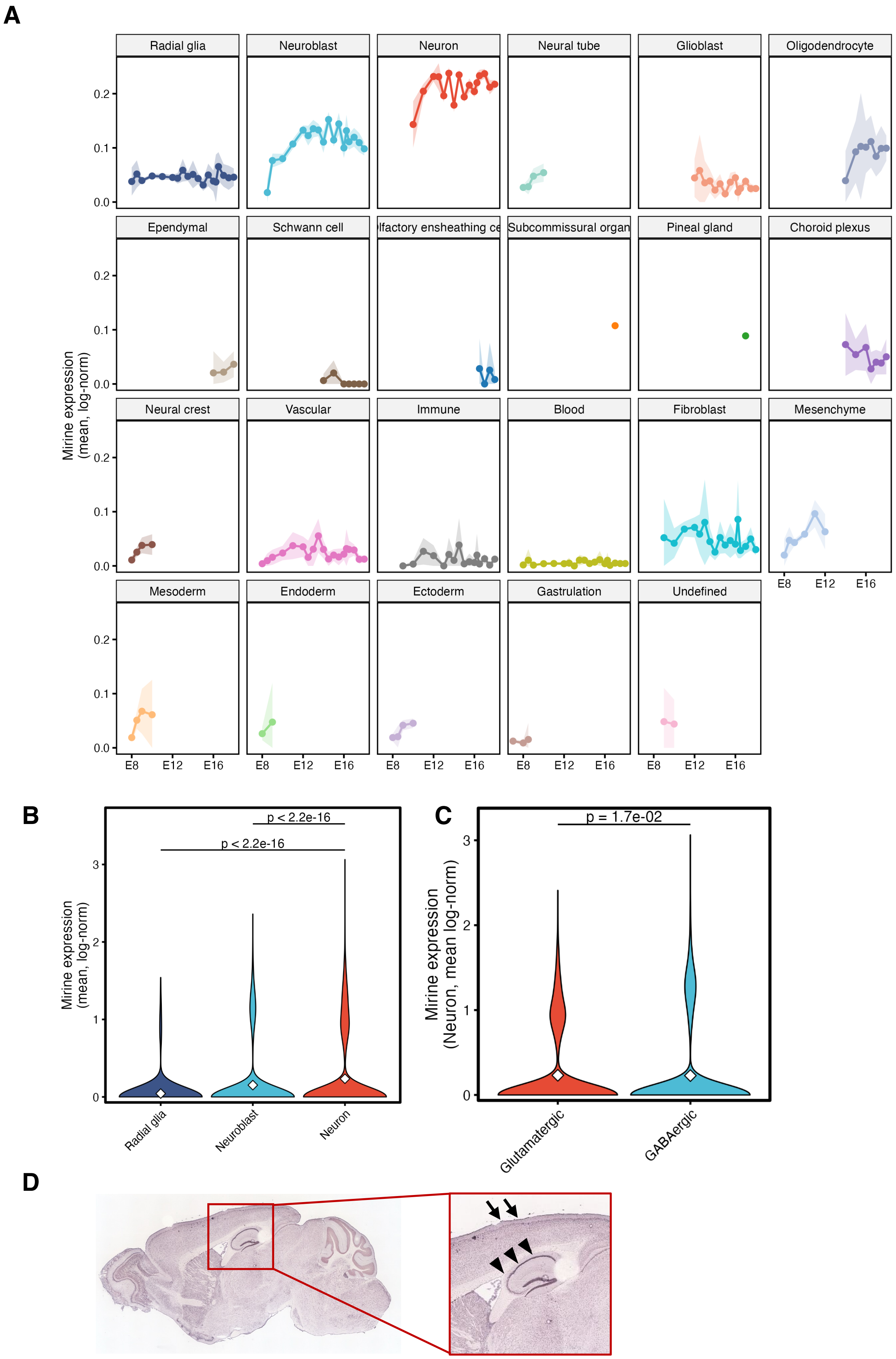

#### Supplementary Figure 4

**A**

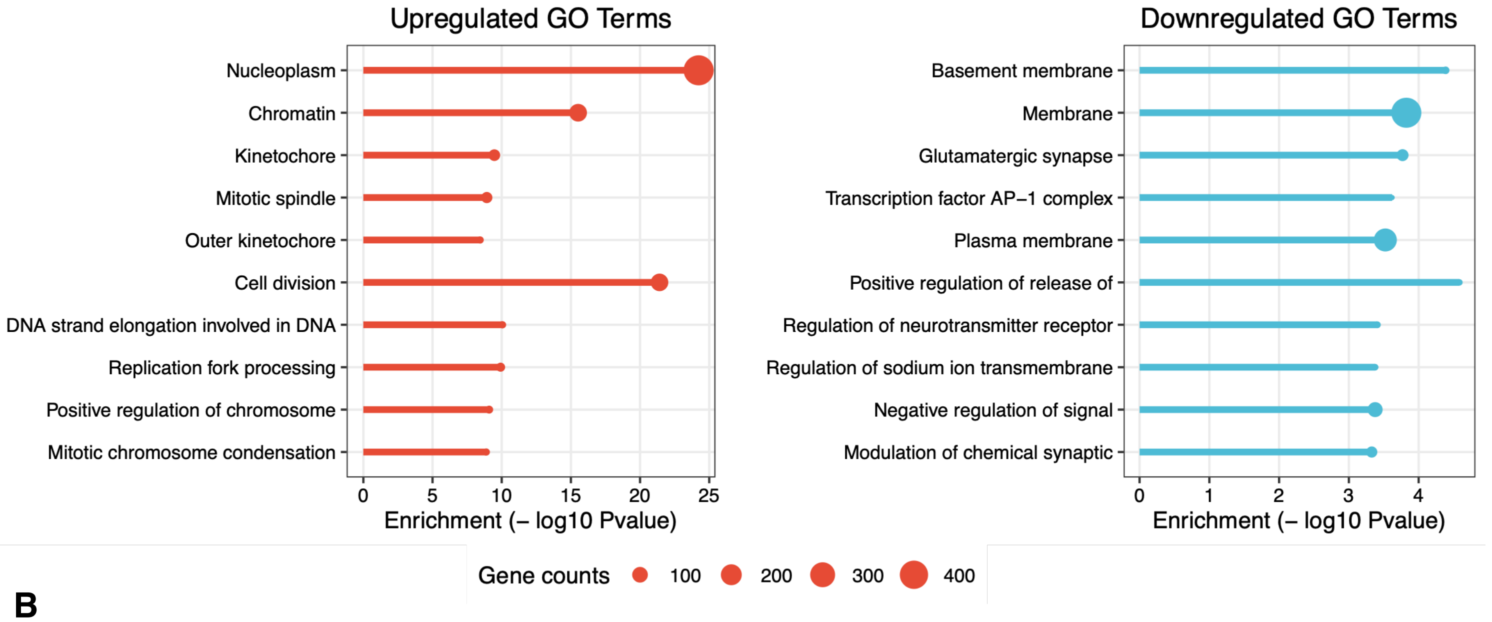


**B**

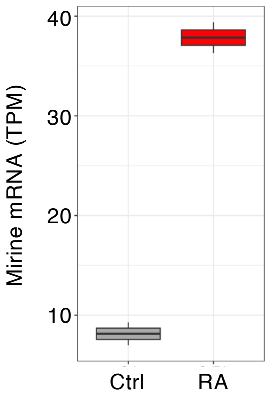

#### Supplementary Figure 5

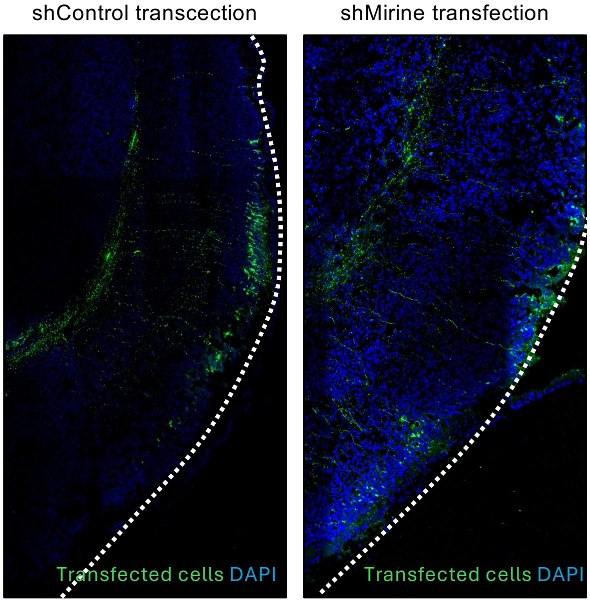

#### Supplementary Figure 6

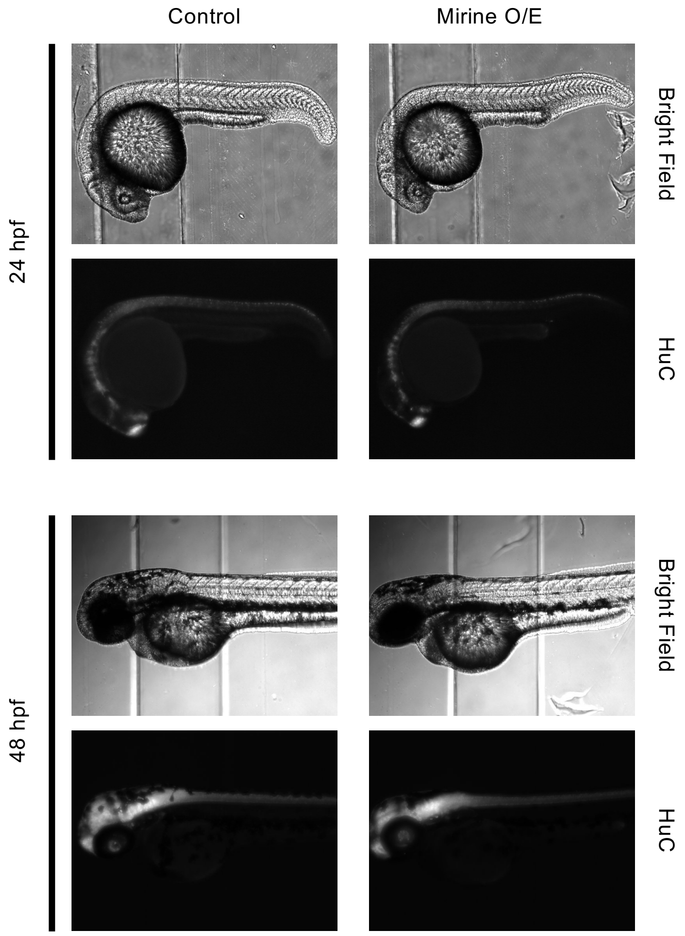

#### Supplementary Figure 7

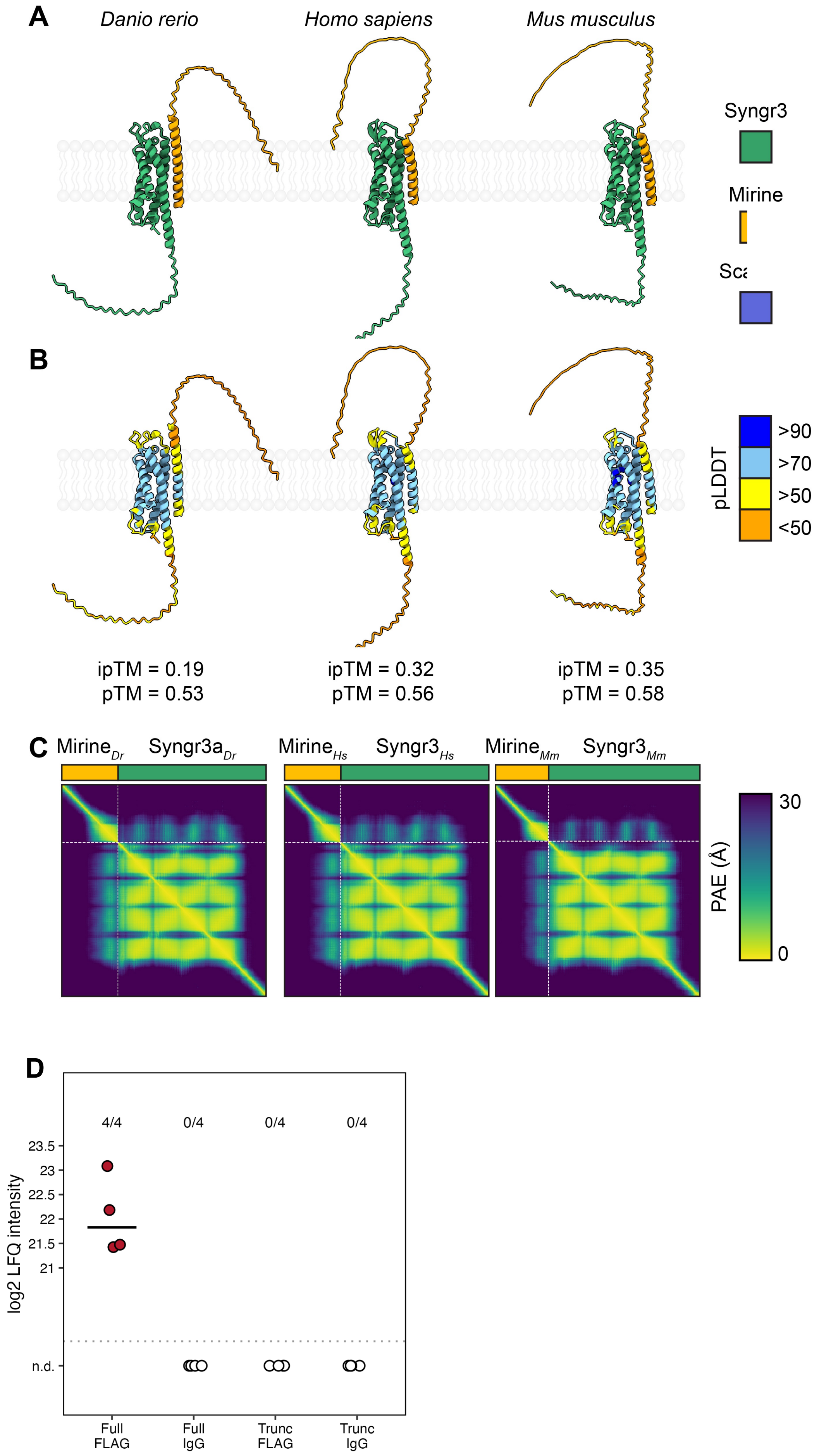
